# Ultra-large targeted DNA integrations in primary human cells

**DOI:** 10.64898/2026.04.09.717505

**Authors:** Courtney Kernick, Lauren Chow, Mikail Alejandro, Ke Li, Maxwell G. Foisey, Xiaoyu Yang, Claire Hilburger, Johnathan Lu, Lujing Wu, Alison McClellan, Oliver Takacsi-Nagy, Rocco Brajenovic, Nicole E. Theberath, Emily Celallos Fuentes, Erin Lin, Austin Hartman, Tina Truong, Jae Hyun Jenny Lee, Luke Workley, Alvin S. Ha, Yongchang Ji, Nicholas Putnam, Nektaria Andronikou, Nujhat Fatima, Max Dotson, Kimberly A. Wong, Christian H. Burns, Floris A. S. Engelhard, Elena Stoyanova, Milica Vukovic, Tom Adie, Wendell A. Lim, Kole Roybal, Katherine Santostefano, Ricardo Almeida, Greg M. Allen, Brian R. Shy, Theodore L. Roth

## Abstract

Genetic <u>engineering</u> experiments and therapies are constrained by the size of DNA integrations into human cell’s genomes. Existing AAV, lentiviral, and non-viral methods rapidly decrease in integration efficiency beyond ∼5kb of sequence. Through systematic evaluation of non-viral DNA template formats, we identified circular ssDNA and dsDNA as capable of mediating >5kb integrations. Large circular DNA delivery efficiency and its impacts on cell viability and payload expression could be significantly improved with small DNA “helper” plasmids, mRNA-encoded nucleases, and sequence design optimizations. Collectively, these modifications enabled ultra-large—up to 10 kb DNA—integrations at >20% efficiency in primary human T cells at the *TRAC* locus and at >60% efficiency in human iPSCs at the *AAVS1* locus. Finally, we demonstrate that GMP clinical-manufactured T cells with ultra-large integrations are functional *in vitro* and *in vivo*. Overall, we identified optimal template architectures, delivery modes, and sequence design rules for ultra-large DNA integrations in both research and clinical settings to accelerate basic genetic research and next-generation cellular therapies.

## MAIN

Viral vectors^1,2^, CRISPR systems^3–5^, and other DNA integration technologies such as recombinases^6–8^ and transposases^9–12^ have enabled manipulation of the human genome to uncover basic biologic mechanisms and therapeutically program cell function^13^. Gene therapy applications based on targeted base-pair changes, larger random viral integrations, or targeted non-viral integrations have been used clinically to correct primary immunodeficiencies and hemoglobinopathies in hematopoietic stem cells (HSCs)^14,15^ and *in vivo* for a variety of monogenic disorders^16,17^. Similarly, cellular therapy applications based on these genetic integration methods, such as CAR T cells, have also been applied to cancer and autoimmune therapies^18,19^. The ability to integrate larger DNA sequences has allowed for greater control over human cell function, such as multi-gene cassettes^20,21^, combinatorial antigen recognition^22^ and synthetic logic circuits with multiple protein products and expression control elements^23^. However, these sophisticated modifications require increasingly large DNA cassettes, which can exceed the capacity of current targeted integration methods in primary human cells^24^, particularly in clinical manufacturing, limiting the complexity of translational genetic engineering. In sum, efficient targeted integrations of large DNA sequences to the human genome remains a foundational technical constraint for both basic research and genetic therapies.

Both viral and non-viral methods have been developed to introduce increasingly large DNA templates into defined human genomic loci. In the 1990s, natural lenti- and retroviruses were engineered to deliver DNA into human patients and subsequent design improvements have enabled routine delivery of ∼5-6 kb of DNA, with the potential for up to ∼8-10kb integrations^25–27^. However, lenti- and retroviral vectors integrate pseudo-randomly, which led to notable deaths in early HSC gene therapy trials for primary immunodeficiencies^28–30^ and, more recently, oncogenic transformation of engineered T cell therapies^31,32^. On the other hand, multi-step editing with stringent selection (such as landing pad installation into a cell line followed by recombinase mediated integration) has enabled extremely large integrations (>50kb)^33–35^, but are largely incompatible with primary human cell applications. Thus, targeted, “single-shot” methods capable of integrating large DNA sequences at high enough efficiencies for clinical application are thus of great interest.

Single-shot, targeted integrations in human cells became more broadly feasible with the advent of CRISPR technologies in the 2010s^3–5^. Delivery of an RNA-guided nuclease into primary human cells along with a DNA template for homology-directed repair (HDR) at the site of the nuclease-induced dsDNA break enables site-specific integration^36^. Various formats of DNA templates have been proposed, but some present important trade-offs: (1) extremely short ssDNA ssODNs^37,38^, (2) efficient but size limited AAV genomes (<4.5kb)^39,40^, (3) readily manufacturable but potentially toxic linear and circular dsDNA templates^21,41^, and (4) challenging to manufacture linear^36,42–44^ or circular ssDNA^45–48^. Yet, regardless of template format, the efficiency and edited cell viability of current single-shot targeted integration methods declines precipitously for integrations above ∼5-6 kb. Non-CRISPR-mediated targeted integration has recently been described using engineering recombinases^49,50^, including simultaneous targeted integration of landing pads^51–53,8^, but efficient integration in primary cells has been limited to ∼1 kb. Further, regardless of the enzyme mediating targeted integration, no comprehensive comparisons of DNA template formats have nominated an optimal format for efficient integrations above ∼5-6 kb.

Diverse approaches to improving cell viability, DNA delivery, editing efficiency, and payload expression when introducing large DNA templates have been described. Optimizations of the DNA template sequence have described in various cell lines and some primary human cell types, such as predicting optimal gRNA sequences for HDR repair^54^, identification of genomic loci most amenable for integrations^21,43^, and homology arm length, symmetry, and 5’ and 3’ end structure^55^. Moreover, small molecule inhibitors of NHEJ^56–58^, MMEJ^59^, and DNA sensing pathways^60^ have been shown to improve editing efficiencies, although these may decrease viability and increase genotoxicity risks^61^, which could potentially limit clinical application. More complex approaches such as cross-linking nucleases to DNA templates^62–65^, using nucleases conjugated to additional components^66^, or expressing additional gene products or endogenous gene knockouts to increase HDR frequency^67–70^ have also been described. And finally, low frequencies of edited cells can be positively or negatively selected^71–73^, although this also potentially limits both clinical and genetic screening applications requiring large numbers of successfully edited cells. Overall, despite these improvements, efficient large, targeted gene integrations above ∼5-6 kb have never been demonstrated in primary human cells^74,21,75^.

Here, we describe the development of Generalized Large-Integrations by DNA Electroporation (GLIDE) editing, a high efficiency method for ultra-large targeted DNA integrations of >10 kb in human cells, including primary human T cells and human iPSCs. Through a systematic evaluation of non-viral DNA template formats, we identified circular ssDNA and circular dsDNA templates as potentially compatible with transgene integrations above 5 kb in length. Inclusion of small “helper” plasmids, use of mRNA encoded nucleases, and a suite of DNA sequence design optimizations drastically improved the viability and efficiency of integrations using circular dsDNA templates. Application of GLIDE-editing in primary human T cells enabled efficient >10kb integrations at the *TRAC* locus. Further, GLIDE-editing’s suite of targeted DNA integration tools proved to be cell type independent, directly enabling efficient > 8 kb integrations at the *AAVS1* locus in multiple human iPSC lines as well. Finally, ultra-large targeted integrations were compatible with GMP clinical manufacturing, and the resulting edited human T cell products showed robust function *in vitro* and *in vivo*.

## RESULTS

### Comparison of DNA template formats for ultra-large knock-ins in primary human T cells

We previously developed several non-viral strategies for targeted insertion of large DNA payloads in primary human T cells using engineered dsDNA or ssDNA templates incorporating truncated Cas9 Target Sites (tCTS, containing a 4 bp mismatch at the 5’ end of the gRNA target sequence to support Cas9 binding without cleavage), or full length Cas9 Target Sites (CTS) to improve efficiency and yield while reducing DNA toxicity^36,76,43^. Additional template formats such as circular ssDNA (cssDNA)^45–48^, traditional circular dsDNA plasmids^77^, and plasmids with minimal backbone (e.g. nanoplasmid or minicircle plasmids)^41,45^ have also shown promise in individual studies. Thus, we initially sought to systematically test a broad range of DNA template formats to identify the most promising formats for further development of larger DNA integrations. Using a previously validated ∼1.6 kb BCMA-CAR construct for targeted knock-in at the *TRAC* locus, we compared linear, circular, viral, ssDNA, and dsDNA template variants (ranging in total size from 2.9 kb for linear templates to 5.6 kb for circular dsDNA plasmid templates), with or without tCTS sites^43,76^ (**Fig. 1a** and **Supplementary Table 1**). Non-viral DNA templates and Cas9 ribonucleoproteins were co-delivered into activated primary human T cells by electroporation as previously described^36,76,43^.

**Figure 1.**
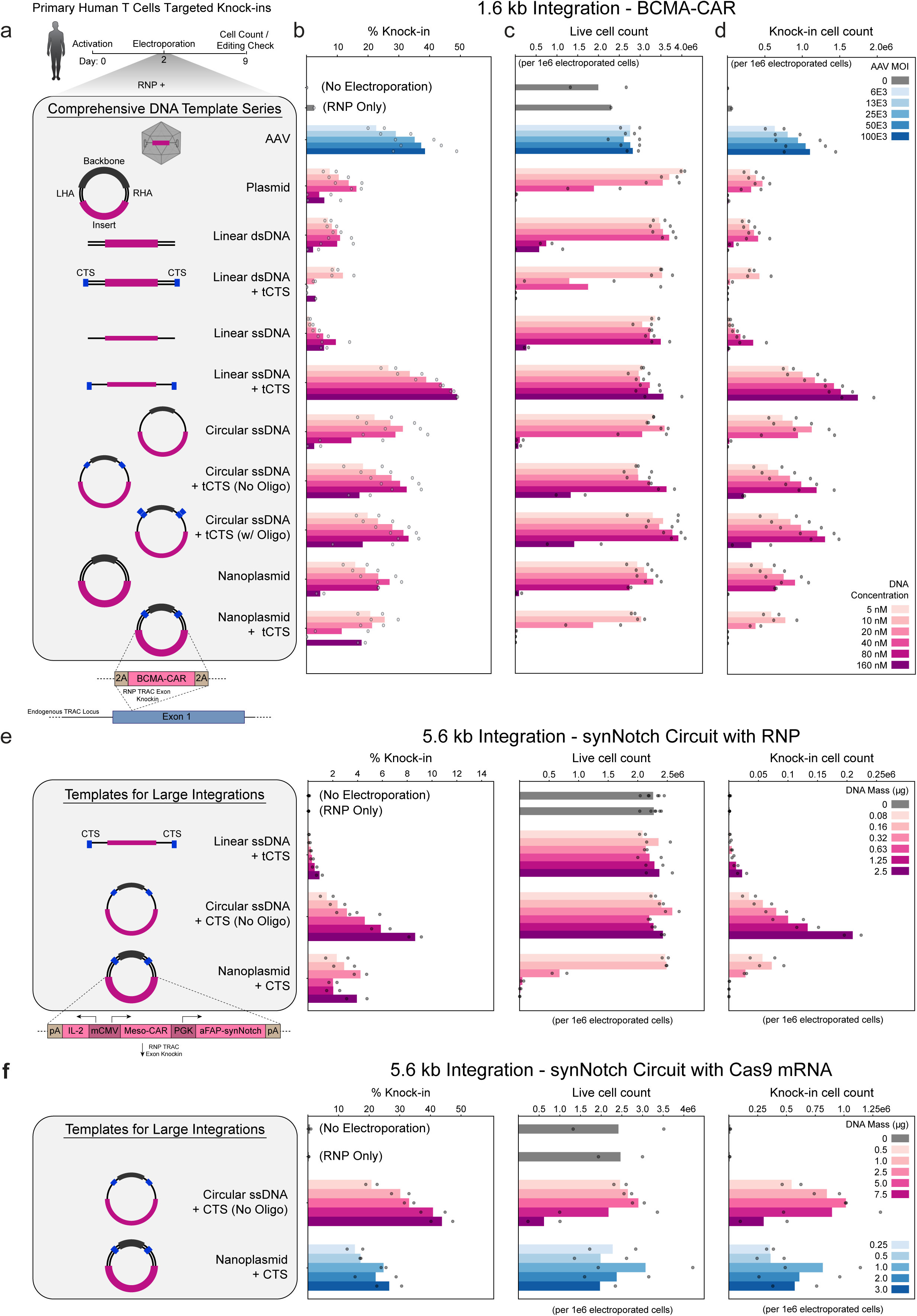
Comprehensive comparison of DNA template types for targeted integration. **(a)** Timeline of T cell knock-in electroporation workflow, knock-in strategy and designs for BCMA-CAR (1.6 kb integration) across the series of DNA HDR template formats tested. **(b)** Comparison of BCMA-CAR HDRTs at concentrations 5nM-160nM or 0-100E3 MOI in terms of knock-in efficiency, **(c)** live cell count per 1e6 edited cells, and **(d)** knock-in cell count per 1e6 edited cells measured 7 days post electroporation. **(e)** Knock-in of a logic-gated synNotch circuit (5.6 kb integration) using linear ssDNA + tCTS, circular cssDNA + CTS, and nanoplasmid + CTS templates with corresponding knock-in efficiency, live cell count and knock-in cell count 7 days post electroporation using Cas9 mRNA. Circular cssDNA was produced and provided by Kano Therapeutics. **(f)** Knock-in strategy and designs for a logic-gated synNotch circuit (5.6 kb integration) at concentrations 5nM-160nM with corresponding knock-in efficiency, live cell count, and knock-in cell count 7 days post electroporation using Cas9 RNP or Cas9 mRNA. Each experiment was performed with T cells from two independent healthy human blood donors represented by individual dots plus mean. CTS, Cas9 target site. RNP, ribonucleoprotein. MOI, multiplicity of infection.

Primary human T cells were evaluated five days after electroporation of these small 1.6 kb integration vectors for knock-in efficiency (**Fig. 1b**), viability (**Fig. 1c**), and knock-in cell yield (**Fig. 1d**). Similar to prior studies, linear ssDNA templates with tCTS sites provided high efficiency and yield of DNA insertion with minimal cellular toxicity, at levels comparable to AAV-encoded repair templates. Templates lacking tCTS showed dramatically reduced performance, confirming that tCTS elements are required for optimal activity with linear ssDNA. In contrast, cssDNA templates showed slightly lower knock-in efficiencies compared to linear ssDNA regardless of tCTS inclusion (**Figure 1a-d**). The cssDNA template was improved by incorporation of a fully gRNA matched CTS (i.e. no 4bp mismatch), which enabled Cas9 cleavage and linearization and resulted in comparable knock-in efficiencies and cell counts achieved with linear tCTS ssDNA (**Extended Data Fig. 1**), similar to improvements in editing efficiency seen with inclusion of a full CTS in circular dsDNA templates^78,79,75^.

Across dsDNA formats, linear dsDNA and standard circular dsDNA plasmids showed increased toxicity and lower knock-in efficiencies relative to optimal ssDNA templates. However, circular dsDNA nanoplasmid templates with a minimal <500 bp backbone significantly improved knock-in rates and reduced cellular toxicity in comparison to alternative dsDNA variants^41^. Evaluation of template delivery and stability over a 7-day time course using matched molar concentrations of each variant revealed substantially higher levels of initial DNA delivery using dsDNA templates, with both linear and circular ssDNA templates also showing shorter half-life within T cells as well (**Extended Data Fig. 2**).

Altogether, ssDNA templates demonstrated the highest knock-in rates and yield for the reference ∼1.6 kb integration BCMA-CAR construct. However, more powerful genetic discovery pipelines and next generation clinical cellular therapies require increasingly large payload sizes to incorporate components such as genetic circuits, safety switches, potency enhancements, and additional modifiers. To evaluate template class performance with larger integration sizes, we tested *TRAC* locus knock-in constructs incorporating an anti-B7H3 CAR + CARD11-PIK3R3 fusion potency enhancement (∼3.5 kb integration) (**Extended Data Fig. 1b**) or a larger logic-gated multi-receptor synNotch circuit (∼5.6 kb integration)^23,80,81^ (**Fig. 1e**). Surprisingly, linear tCTS ssDNA templates showed a rapid drop in performance, decreasing from 40-50% knock-in with the 1.6 kb integration to less than 2-3% for 3.5 and 5.6 kb integrations (**Fig. 1e** and **Extended Data Fig. 1b**). In contrast, circular dsDNA nanoplasmid CTS templates supported approximately 20-30% knock-in at 3.5 kb integration and approximately 5% at 5.6 kb integration, albeit with significant dose-dependent cellular toxicity. cssDNA templates demonstrated further benefit, achieving ∼5-10% knock-in for the 5.6 kb integration with minimal toxicity (**Fig. 1e**).

This comprehensive comparison of DNA template types for targeted DNA integrations revealed a clear size limit for linear ssDNA templates and nominated circular DNA formats, both circular ssDNA and circular dsDNA plasmids and nanoplasmids, as preferred strategies for ultra-large payload delivery. Indeed, circular dsDNA plasmids of up to ∼14 kb in total template size could be delivered into primary T cells by electroporation (**Extended Data Fig. 3**), albeit with a concomitant template size-dependent viability decrease. Further, we unexpectedly observed that at the largest 5.6 kb integration size tested, switching from pre-assembled Cas9 ribonucleoproteins (RNPs) to Cas9 mRNA reduced the cellular toxicity and dramatically improved knock-in yield for both formats, achieving ∼20-30% knock-in for the circular dsDNA nanoplasmid template and ∼30-40% knock-in with cssDNA (**Fig. 1f** and **Extended Data Fig. 4**). These data highlighted that ultra-large DNA templates, especially circular DNA formats, could be delivered into primary human cells and that potential optimization routes may be available to achieve workable editing efficiencies and cell viabilities.

### Optimized delivery of ultra-large DNA templates into human T cells

We and others have previously identified that delivery of both large positively charged macromolecules (e.g. RNP complexes) and large negatively charged macromolecules (e.g. long DNA sequences with their negatively charged phosphate backbone) by electroporation can be facilitated by co-delivery with long negatively charged polymers, such as shorter DNA sequences^82,83^ or poly-glutamic acid (PGA)^76^. Additionally, we observed that co-delivery of large DNA templates with an mRNA-encoded nuclease, also a long negatively charged polymer, improved cell viability and editing efficiency relative to co-delivery with a positively charged RNP (**Fig. 1f**). Accordingly, we (1) sought to explore how polymer co-delivery impacted the viability and delivery efficiency of larger DNA templates and (2) test whether the polymer-mediated enhancement differs between RNP- and mRNA-based editing contexts.

We initially examined using a small circular dsDNA “helper” plasmid as a co-delivery polymer for comparatively larger circular dsDNA plasmids, each bearing an episomal GFP expression cassette to allow assessment of DNA delivery efficiency and viability post-electroporation (**Fig. 2a**). In primary T cells, we observed that increasing sizes of the circular dsDNA template (e.g. an HDRT) caused step-wise decreases in viability relative to electroporation only, but inclusion of an additional small helper plasmid (∼4 kb total size, with no homology to DNA HDRT) as a co-delivery polymer rescued this viability loss, with increasing effect for larger circular dsDNA plasmid templates (**Fig. 2b**). This improved viability was also associated with improved delivery of the template plasmid, again an effect that appeared to be size dependent **(Fig. 2c).** Optimization of the co-delivery helper plasmid size, concentration, template to helper ratios, volumes, and mass ultimately yielded a standardized protocol for inclusion of small helper plasmids that increased editing efficiency and viability for *TRAC* locus knock-ins in primary T cells across >90% of HDR templates tested, especially for large integrations above >5 kb (**Extended Data Fig. 5**).

**Figure 2.**
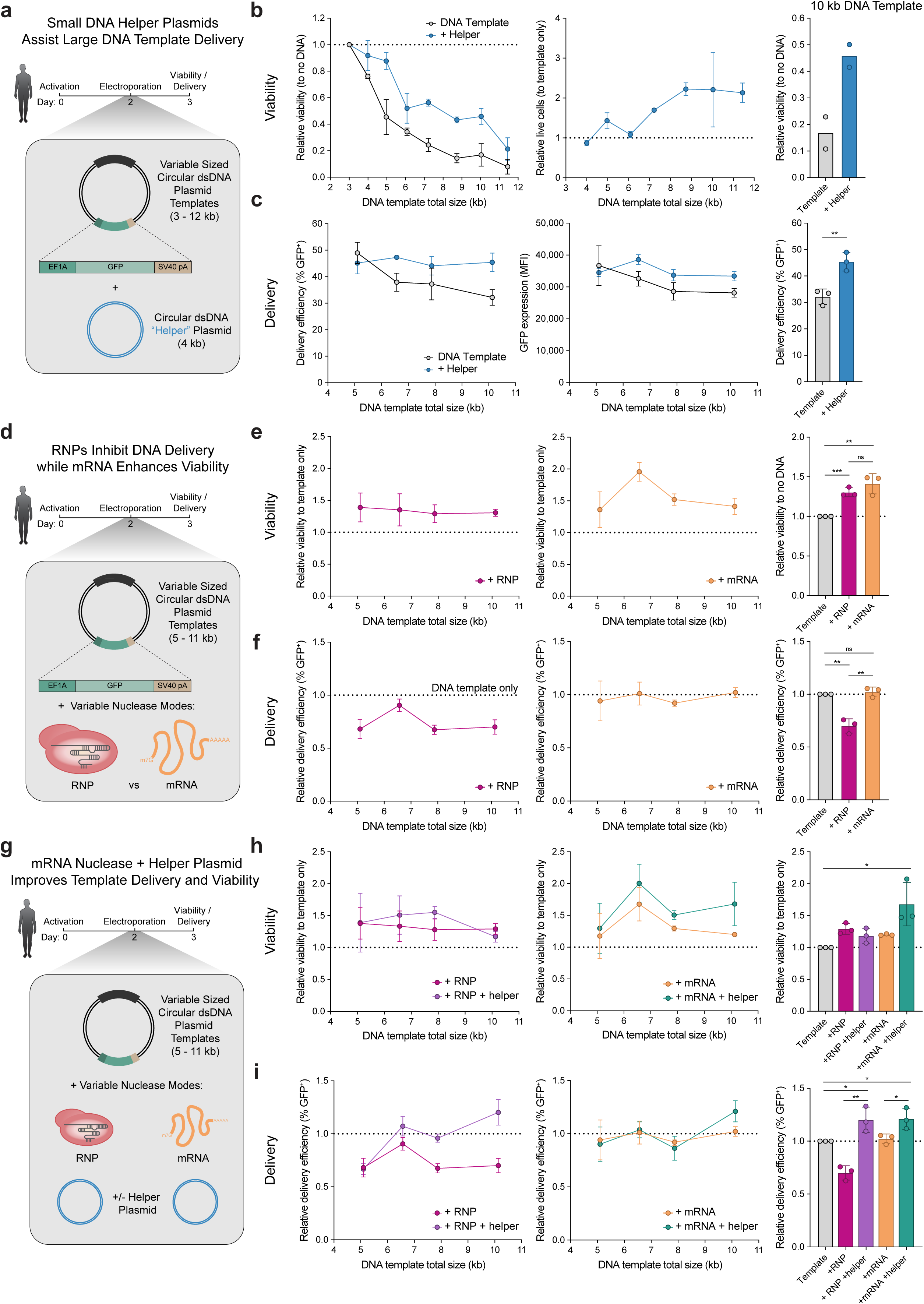
Small DNA Helper plasmids and mRNA templates improve delivery and reduce toxicity of large DNA sequences. **(a)** Electroporation of primary human T cells with episomal GFP expression plasmids of increasing size (DNA templates) and helper plasmid. **(b)** Live cell counts 24 hours post-electroporation of cells receiving no DNA, DNA template alone, or DNA template with helper plasmid (*n* = 2 donors). **(c)** DNA template delivery efficiency 24 hours post-electroporation, measured by GFP^+^ cell frequency and mean fluorescence intensity (MFI) of GFP^+^ cells (*n* = 3 donors, mean ± s.d.). \*\**P* < 0.01, two-sided unpaired t test. **(d)** Electroporation of primary human T cells with DNA templates and Cas9 RNP or Cas12a mRNA. **(e,f)** Live cell counts **(e)** and delivery efficiency **(f)** 24 hours post-electroporation, compared to DNA template-only controls (*n* = 3 donors, mean ± s.d.). ns, \*\**P* < 0.01, \*\*\**P* < 0.001, two-sided unpaired t test. **(g)** Electroporation of primary human T cells with DNA templates, Cas9 RNP or Cas12a mRNA, and helper plasmid. (**h,i)** Live cell counts **(h)** and delivery efficiency (**i)** 24 hours post-electroporation compared to DNA template-only controls (*n* = 3 donors, mean ± s.d.). ns, \**P* < 0.05, \*\**P* < 0.01, \*\*\**P* < 0.001, two-sided unpaired t test.

We then asked whether the negatively-charged mRNA improved editing in part via a co-delivery effect and if this could be further optimized. While inclusion of either RNPs or mRNA nucleases during electroporation with a DNA template improved viability across template sizes (a finding we previously reported for RNPs^36^), RNPs appeared to do so by decreasing the efficiency of DNA template delivery (**Fig. 2e,f**). In contrast, mRNA improved viability with no impact on delivery efficiency (**Fig. 2b-f)**. Combining large DNA template electroporation with co-delivery of RNP + helper plasmid or mRNA + helper plasmid revealed that helper plasmids improved viability for both RNP and mRNA conditions, whereas improved delivery efficiency was most pronounced with RNPs at large DNA template sizes (**Fig. 2g-i**). Overall, co-delivery of mRNA-encoded nucleases and small helper plasmids with a large (∼10kb total template size) circular dsDNA template improved cell viability by >50% and DNA template delivery by >20%, with minimal additional cost or complexity.

### Miniaturization and optimization of ultra-large DNA knock-in sequences

Having demonstrated that significantly larger DNA templates could be introduced into primary human T cells through co-delivery with mRNA-encoded nucleases and small helper plasmids, we next sought to maximize the amount of the delivered DNA template that could be genomically integrated into a target locus (**Fig. 3a**). Even with helper plasmids and mRNA nucleases, the largest circular dsDNA plasmid we could deliver was ∼14 kb (**Fig. 2b** and **Extended Data Fig. 3**). Thus, we sought to minimize DNA template components not part of the integrated sequence, specifically the “targeting” sequences, such as homology arms, and the plasmid backbone itself. Typically, long homology arms (∼1 kb) are included on either side of an integrated sequence to mediate HDR at the site of nuclease-mediated DSB^21,41^. Homology-independent targeted integrations (HITI)^84,85^ and homology mediated end joining (HMEJ) / microhomology-mediated (MMEJ) integrations^78,74^ both require significantly shorter (0-50 bp) homology arms than HDR (**Fig. 3b**). For small ∼2.6 kb integrations at the *TRAC* locus, we found that MMEJ format templates with ∼50 bp homology arms flanked by CTSs for linearization of the circular dsDNA template was as efficient as canonical HDR with ∼1 kb homology arms. However, for larger total DNA templates sizes of greater than 10 kb, MMEJ format templates proved to be significantly more efficient, achieving ∼16% knock-in compared to ∼6.5% with HDR templates. We note that the alternative short homology arm-based repair pathway, HITI, was not efficient at the tested loci, which may be due to the specific gRNA cut site, as previously described^84^ (**Fig. 3c**). Directionality of the CTS sequence in MMEJ format templates (either “In” with the PAM 5’-3’ oriented towards the integrated sequence, or “Out” facing away) did not appear to impact editing efficiency^76^ (**Fig. 3c**). MMEJ format templates, which inherently contain intact Cas9/12a Targeting Sequences also showed lower levels of off-target integrations compared to non-CTS containing DNA templates (**Extended Data Fig. 6**). Miniaturization of plasmid backbone sequences similarly yielded increases in editing efficiency (**Fig. 3d,e**), including a minimal 1.6 kb plasmid backbone (pUCmu)^86^, providing ∼50% of the total size reduction of nanoplasmids or minicircles^87–89^ without requiring specialized bacterial strains or complex recombination/excision, respectively.

**Figure 3.**
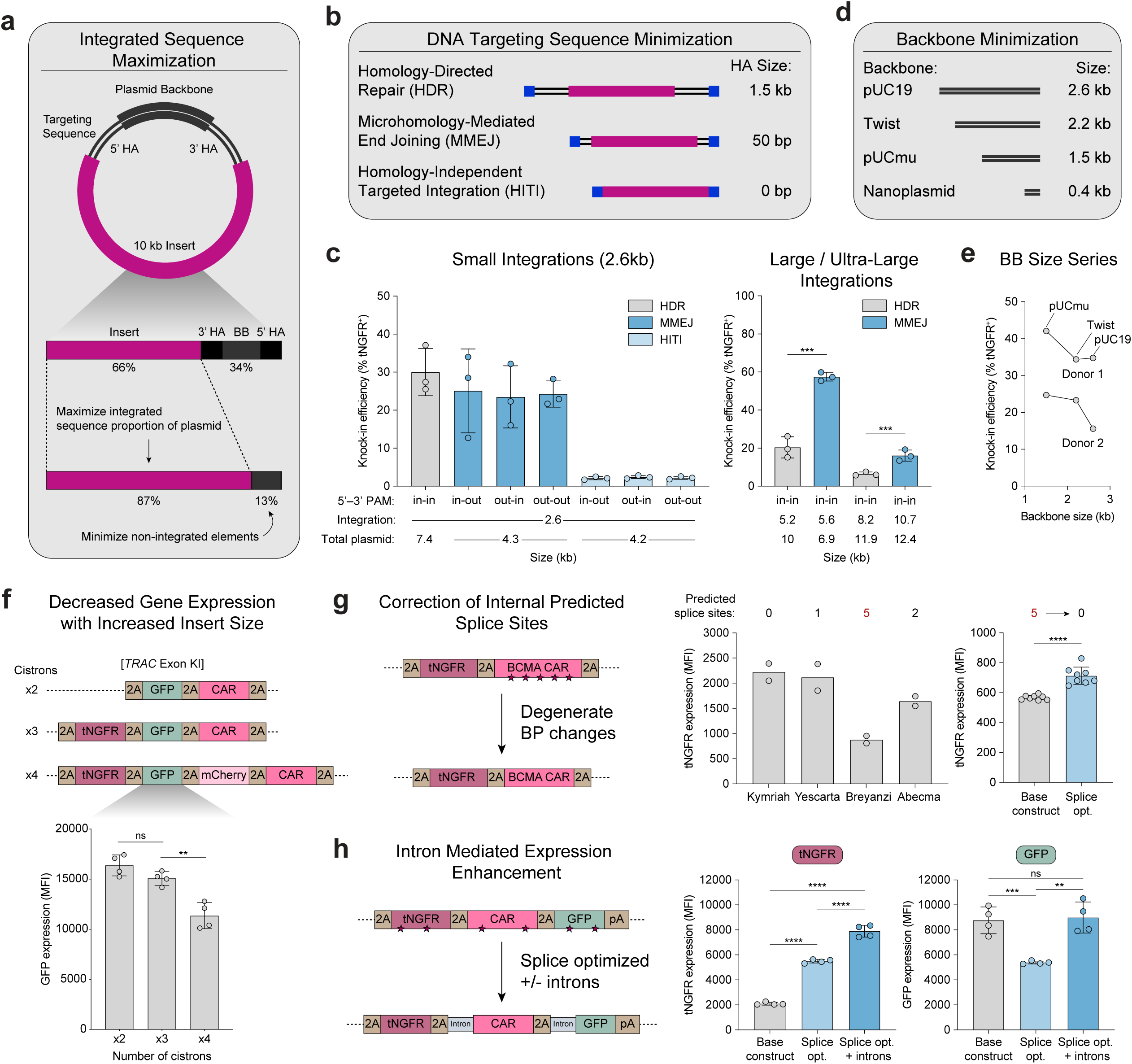
Maximization and optimization of integrated sequences with large DNA templates. **(a)** Strategies to maximize integrated sequence size within practical DNA template size constraints, enabling larger genomic integrations in primary cells. **(b)** Minimizing targeting sequence length by leveraging alternative DNA repair pathways (MMEJ or HITI). **(c)** Knock-in efficiency across HDR, MMEJ and HITI templates, measured in edited (tNGFR^+^) cells 11 days post-electroporation (*n* = 3 donors, mean ± s.d.). \*\*\**P* < 0.001, two-sided unpaired t test. **(d)** Minimizing plasmid backbone length. **(e)** Knock-in efficiency of HDR templates (4 kb integration) with varying backbone sizes, measured in edited (tNGFR^+^) cells 5 days post-electroporation (*n* = 2). **(f)** Expression from multicistronic cassettes of increasing size (2.5 kb, 3.4 kb, and 4.1 kb integrations) integrated at the *TRAC* locus. MFI of edited (GFP^+^) cells 5 days post-electroporation (*n* = 2 donors, 2 technical replicates, mean ± s.d.). ns, \*\**P* < 0.01, two-sided unpaired t test. **(g)** Expression of five FDA-approved CARs using tNGFR as a reporter. MFI in edited (tNGFR^+^) cells (n = 2). Breyanzi CAR expression before and after splice site removal (*n* = 2 donors, 4 technical replicates, mean ± s.d.). ns, \*\*\*\**P* < 0.0001, two-sided unpaired t test. **(h)** Expression of a bicistronic tNGFR-GFP cassette (unmodified, splice site-corrected, or splice site-corrected with short introns) integrated at the *TRAC* locus. MFI in edited (tNGFR^+^) cells 7 days post-electroporation (*n* = 2 donors, 2 technical replicates, mean ± s.d.). ns, \*\**P* < 0.01, \*\*\**P* < 0.001, \*\*\*\**P* < 0.0001, two-sided unpaired t test.

Our tests of large multi-cistronic cassettes revealed expression liabilities that were not apparent in smaller multi-cistronic cassettes^36,76,43,20^. Namely, as the number of genes within a multi-cistronic cassette increased, the expression level of individual genes decreased, as reported previously^21^ (**Fig. 3f**). While we did not comprehensively examine approaches to rescue size dependent deceases in expression efficiency with ultra-large DNA integration sizes, we did identify simple efforts that were not previously necessary for smaller one to two gene integrations. We observed differences in expression between bi-cistronic cassettes with one identical reporter gene and one variable chimeric antigen receptor (CAR) gene that was potentially attributable to the frequency of predicted splice sites in the CAR **(Fig. 3g)**. Further, inclusion of short introns^90,91^ between distinct genes in a tri-cistronic cassette also rescued expression levels (**Fig. 3h**). Both systematic mutation of predicted consensus splice donor and acceptor sequences (using degenerate codons), as well as intronization —the inclusion of short intronic sequences mimic intron-exon architecture of natural genes—were simple DNA modifications that were able to partially to rescue expression levels for large multi-cistronic sequences (**Fig. 3g,h**).

### Generalized large integrations by DNA electroporation (GLIDE) editing across human cell types

We next sought to combine the improvements we identified in ultra-large DNA template format, delivery, viability, and sequence design into a single method: Generalized Large Integrations by DNA Electroporation, or GLIDE-editing (**Fig. 4a**). GLIDE-editing in primary human T cells resulted in integration of a 10.7 kb sequence containing a seven-gene cassette at *TRAC* using a 12.4 kb total sized circular dsDNA plasmid template and represented a ∼50% increase in viability, ∼4-fold increase in editing efficiency, and a ∼3-12-fold increase in total edited cells across six human donors compared to previous methods^21^ (**Fig 4b-d**). In addition to improving edited cell yields by almost an order of magnitude, GLIDE-editing optimized ultra-large integration templates were also more efficiently expressed, averaging ∼10-15% improved expression over prior optimized sequences (**Fig. 4e**). As all initial testing and optimization work to develop GLIDE-editing was done in primary human T cells, we wondered whether GLIDE-editing may be generalizable to other human cell types to broadly enable large DNA integrations without additional technical development. We designed a 8.2 kb integration template (within a 9.9 kb circular dsDNA plasmid) for the *AAVS1* safe harbor locus, which is accessible across cell types^92^ (compared to T cell-specific *TRAC* integration), and tested whether GLIDE-editing could be applied to human iPSCs (**Fig. 4f**). Indeed, with only optimization of electroporation pulse parameters (**Extended Data Fig. 7**) >60% integration in KOLF2.1J cells, a widely used standard human iPSC line for genetic engineering and differentiation work^93^. Moreover, ∼20% efficiency was attained in the SCTi004 iPSC line (**Fig. 4g,h**). GLIDE-editing thus appeared to enable off-the-shelf large DNA integrations across multiple human cell types with minimal cell-type specific optimization.

**Figure 4.**
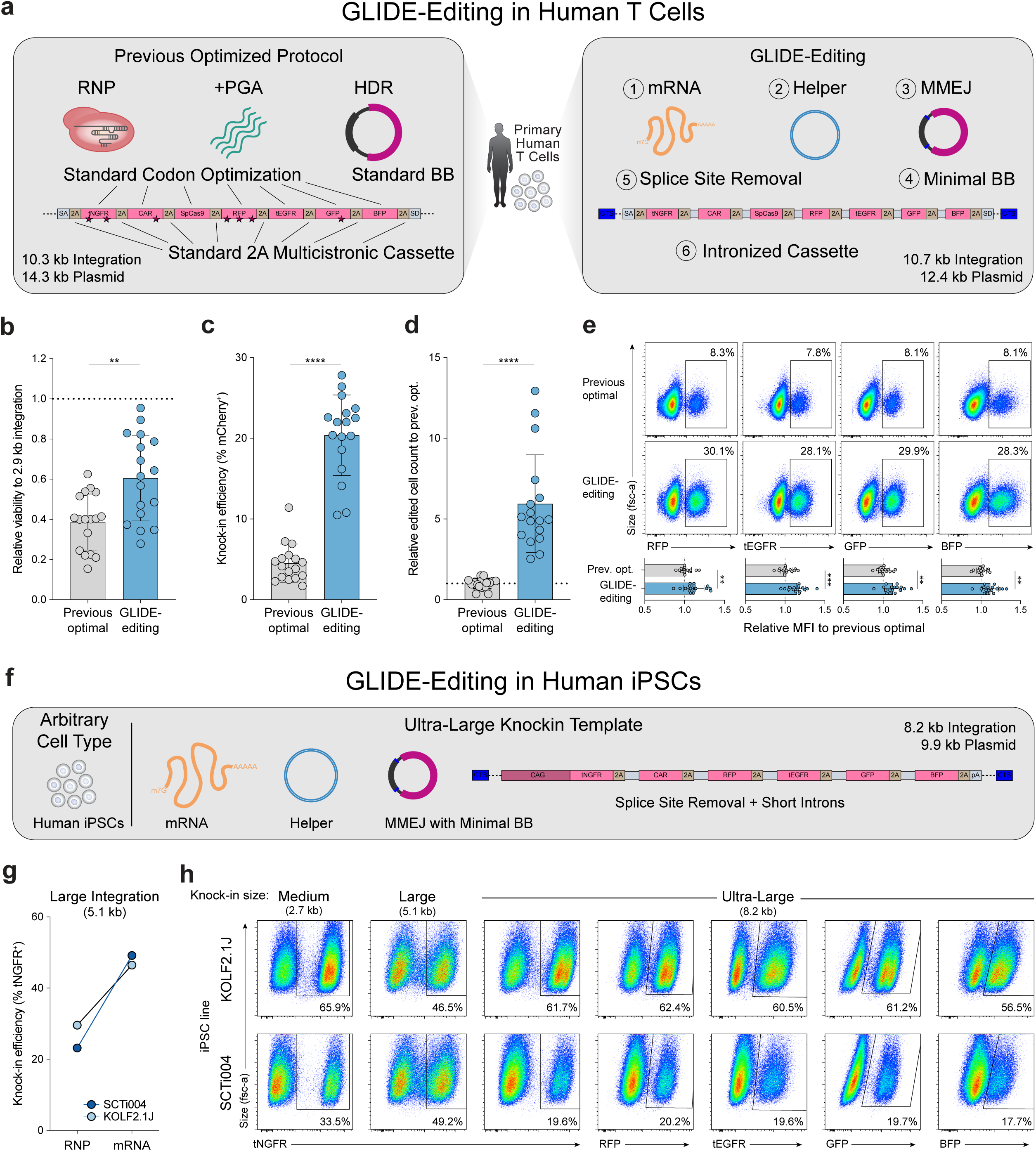
Generalized large integrations by DNA electroporation across multiple human cell types. **(a)** The GLIDE-editing method, incorporating optimizations to improve cell viability, DNA template delivery, and knock-in efficiency for ultra-large integrations in primary human T cells. **(b)** Live cell counts 24 hours post-electroporation compared to a small HDR template control (7.6 kb total template, 2.9 kb integration) (*n* = 6 donors, 2-3 technical replicates, mean ± s.d.). \*\**P* < 0.01, two-sided unpaired t test. **(c-e)** Knock-in efficiency **(c)**, total number of edited cells **(d)**, and gene expression **(e)** with GLIDE-Editing compared to the previous optimized protocol. Knock-in and MFI in edited (RFP^+^) cells 7 days post-electroporation (*n* = 6 donors, 2-3 technical replicates, mean ± s.d.). \*\**P* < 0.01, \*\*\**P* < 0.001, \*\*\*\**P* < 0.0001, two-sided unpaired t test. **(f)** Application of GLIDE-editing to induced pluripotent stem cells (iPSCs). **(g)** Knock-in efficiency (5.1 kb integration) with Cas9 mRNA versus RNP in two iPSC lines. Knock-in measured in edited cells (tNGFR^+^) 6 days post-electroporation. **(h)** Representative flow cytometry plots showing knock-in efficiency across integration sizes in two iPSC lines 6 days post-electroporation.

### GMP-compatible T cell engineering with ultra-large therapeutic constructs

Finally, we sought to demonstrate that ultra-large DNA integrations are compatible with clinical-grade workflows using circular dsDNA templates. As GMP-grade clinical cell manufacturing requires larger volume electroporation instruments compared to research-grade production, we identified electroporation pulse parameters on the CTS Xenon Electroporation System using a minimal “Design of Experiment” strategy (**Extended Data Fig. 8**). Initial testing with a 5.6 kb synNotch circuit integration circular dsDNA nanoplasmid template (7.3 kb total template size) showed an expected inverse relationship between knock-in efficiency and cellular toxicity, although total yields of viable knock-in cells remained high across the tested range (**Fig. 5a**), demonstrating that large DNA templates could be delivered into human T cells at clinical scale.

**Figure 5.**
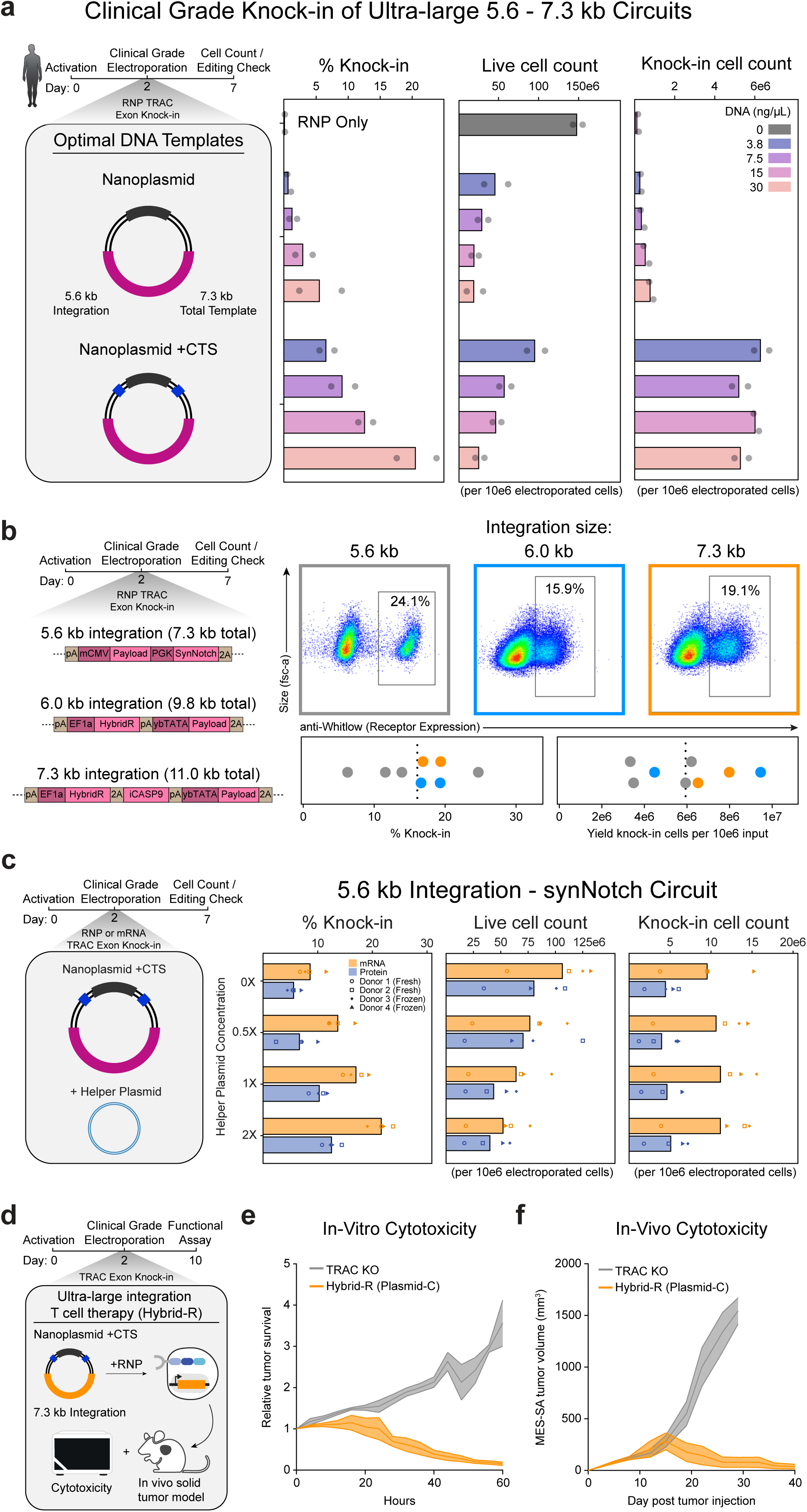
Ultra-large targeted knockins in clinical manufacturing settings. **(a)** Diagram of clinical-scale T cell engineering workflow using Cas9 RNP and a nanoplasmid DNA template encoding a SynNotch circuit (5.6 kb integration). Nanoplasmid templates with a CTS significantly increased knock-in efficiency without additional toxicity. CTS-nanoplasmid showed inverse relationship between knock-in efficiency and cellular toxicity across tested template doses, yielding similar viable knock-in cells. Knock-in percentage, live cell count, and total knock-in cell count measured 5 days poste electroporation. **(b)** Application of the optimized Xenon electroporation protocol to ultra-large nanoplasmid templates with various integration sizes. Representative flow plots and aggregated data showing knock-in efficiencies and yield of viable knock-in cells across constructs. Knock-in percentage and total knock-in cell count measured 5 days poste electroporation. **(c)** Evaluation of ultra-large template integration using the MaxCyte GTx electroporation system. Cas9 was delivered through electroporation in the form of RNP or mRNA, together with a 5.6kb integration (7.3kb total size) construct in the presence and absence of helper plasmid in fresh and frozen donor T cells. Cas9 mRNA increased knock-in efficiency without reducing cellular yield or viability. Inclusion of helper plasmid further increased knock-in efficiency in a dose-dependent manner but was associated with increased cellular toxicity, resulting in stable overall yields of knock-in cells. Knock-in percentage, live cell count, and total knock-in cell count measured 5 days poste electroporation. **(d)** Diagram showing targeted insertion of a 7.3 kb integration (11.0 kb total template size) next-generation anit-B7-H3 Hybrid-R CAR circuit at the TRAC locus using clinical-grade engineering workflow. Engineered T cells were subjected to in vitro and in vivo functional assessment. **(e)** Relative tumor survival (target cell lysis) by anti-B7-H3 Hybrid-Rs targeting B7-H3+ MES-SA mKate-NLS cells as measured by Incucyte live-cell imaging. Effector to target ratio is 1:4. Data are total area normalized to first timepoint (n=3 technical replicates). **(f)** Tumor volumes from B7-H3+ MES-SA tumor-bearing NSG animals treated with 2.0e6 engineered Hybrid-R T cells 5 days after tumor inoculation. n=5, Mean +/- SEM depicted.

We next applied the optimized clinical-scale electroporation protocol to circular dsDNA plasmid templates with up to 7.3 kb integration sizes (11 kb total template size) (**Fig. 5b**). Knock-in efficiencies averaged ∼15% (6-25%) across constructs and produced average yields of roughly 50 million viable knock-in cells from 100 million input T cells in a 1 mL electroporation reaction (**Fig. 5b**). Given the 1 to 25 mL operating range of the Xenon platform, this process is compatible with the generation of hundreds of millions to billions of engineered T cells, well in excess of dose requirements for human patients^43^.

We next evaluated performance using the Maxcyte GTx system, which is an alternative GMP-compatible platform providing comparable electroporation performance across large and small volumes^43^, enabling efficient comparison and optimization across many conditions. In agreement with research-scale experiments using GLIDE-editing, we found that delivery of Cas9 mRNA increased knock-in efficiency for the large 5.6 kb synNotch integration circular dsDNA template (7.5 kb total) by ∼1.5 to 2-fold across all conditions without impact on cellular yield or viability (**Fig. 5c**). Inclusion of helper plasmid similarly increased knock-in efficiency in a dose-dependent manner, albeit with increased cellular toxicity, leading to consistent yield of knock-in cells using baseline electroporation parameters. Performance was broadly similar for fresh and cryopreserved starting material with notable donor variability (**Fig. 5c**).

Finally, we functionally tested human T cells generated by ultra-large DNA integrations at clinical-scale. Gene circuit-regulated CAR T cells that include the inducible caspase 9 (iCasp9) suicide switch^94^ have previously been too large for clinical-grade manufacturing. Using GLIDE-editing, a 7.5 kb integration circular dsDNA template (10 kb total template size) encoding a SNIPR → CAR circuit with an iCasp9 suicide switch was inserted at the *TRAC* locus and exhibited robust and high fidelity SNIPR antigen triggered CAR induction, with selective cytotoxicity of SNIPR antigen positive and CAR antigen positive tumors (**Extended Data Fig 9**). We subsequently performed clinical-scale production with a second previously non-feasible ultra-large DNA integration template— *TRAC* integration of a Hybrid-R circuit^24^ for CAR activation and induction of CARD11-PIK3R3 payload^81^ (7.3 kb integration, 11 kb total template)— which mediated robust cytotoxicity *in vitro* and *in vivo* tumor-clearance at low therapeutic doses (**Fig. 5d-f**). In sum, these studies demonstrate clinically compatible strategies for targeted non-viral insertion of ultra-large constructs, with efficiencies, yields, and function appropriate for clinical use.

## DISCUSSION

Numerous methods to deliver DNA sequences into human cells and integrate those sequences at desired sites within the human genome have been developed and applied. The size of DNA sequence that can be efficiently integrated is a limiting constraint on research and clinical applications for cellular therapies. Within transformed human cell lines or human iPSC capable of stringent selection of cells over the course of weeks, targeted transgene integrations of over 50 kb have been described by clonally isolating extremely rare correct integration events, in particular using recombinase + landing pad-based approaches^34^. However, among “single-shot” gene editing methods potentially compatible with larger scale research or clinical applications in primary human cells, similar targeted recombinase-based approaches have been limited to at most 1 to 2 kb integrations^50–53,8^, AAV-based methods are limited by viral capsid geometry to <4.5 kb, and efficient non-viral targeted integrations have not been reported above ∼5 kb^21^. We describe the development and application of GLIDE-editing, a non-viral method for efficient “one-shot” ultra-large targeted gene integrations in primary human cells. In particular, GLIDE-editing showed efficient >10kb integrations in primary human T cells, efficient ultra-large integrations in iPSCs with minimal optimization and was compatible with clinical grade T cell manufacturing methods.

Numerous formats of DNA templates have been described for use in gene editing applications in human cells, such as linear dsDNA, linear ssDNA, circular dsDNA, and circular ssDNA templates. We report the first comprehensive head-to-head evaluation of each of these template classes containing the same standardized gene integration and characterize the delivery, viability, and editing efficiency in primary human cells (**Fig. 6**). Single-stranded DNA templates, both linear and circular, showed minimal toxicity albeit with decreased delivery efficiency and do require more complex manufacturing. Surprisingly, linear ssDNA showed a marked dropoff in editing efficiency above ∼1.5 kb DNA integrations, while circular ssDNA and circular dsDNA templates were capable of efficient ultra-large >5kb integrations. Toxicity limited efficient linear dsDNA use above ∼1-2 kb integrations, but circular dsDNA templates showed a balance of properties for gene editing usage, with moderate toxicity, greater delivery efficiency, and ease of production at both research and clinical scale. Our comprehensive comparison of the toxicity, delivery efficiency, stability, and editing efficiency across available DNA template classes may be of use beyond targeted HDR/MMEJ DNA integrations to any genetic engineering methodology involving *ex vivo* delivery of DNA into primary cells, such as for episomal expression, transposon-based integrations, and recombinase-based integrations. In particular, circular dsDNA plasmids, the template class we found the most readily applicable for ultra-large sequence delivery into primary cells by electroporation, appeared to have a limiting size cutoff of ∼13-14 kb total sequence length, highlighting that ∼10-12kb integrated sequence lengths may represent a fundamental maximal size in primary human T cells using current *in vitro* DNA delivery methods such as electroporation.

**Figure 6.**
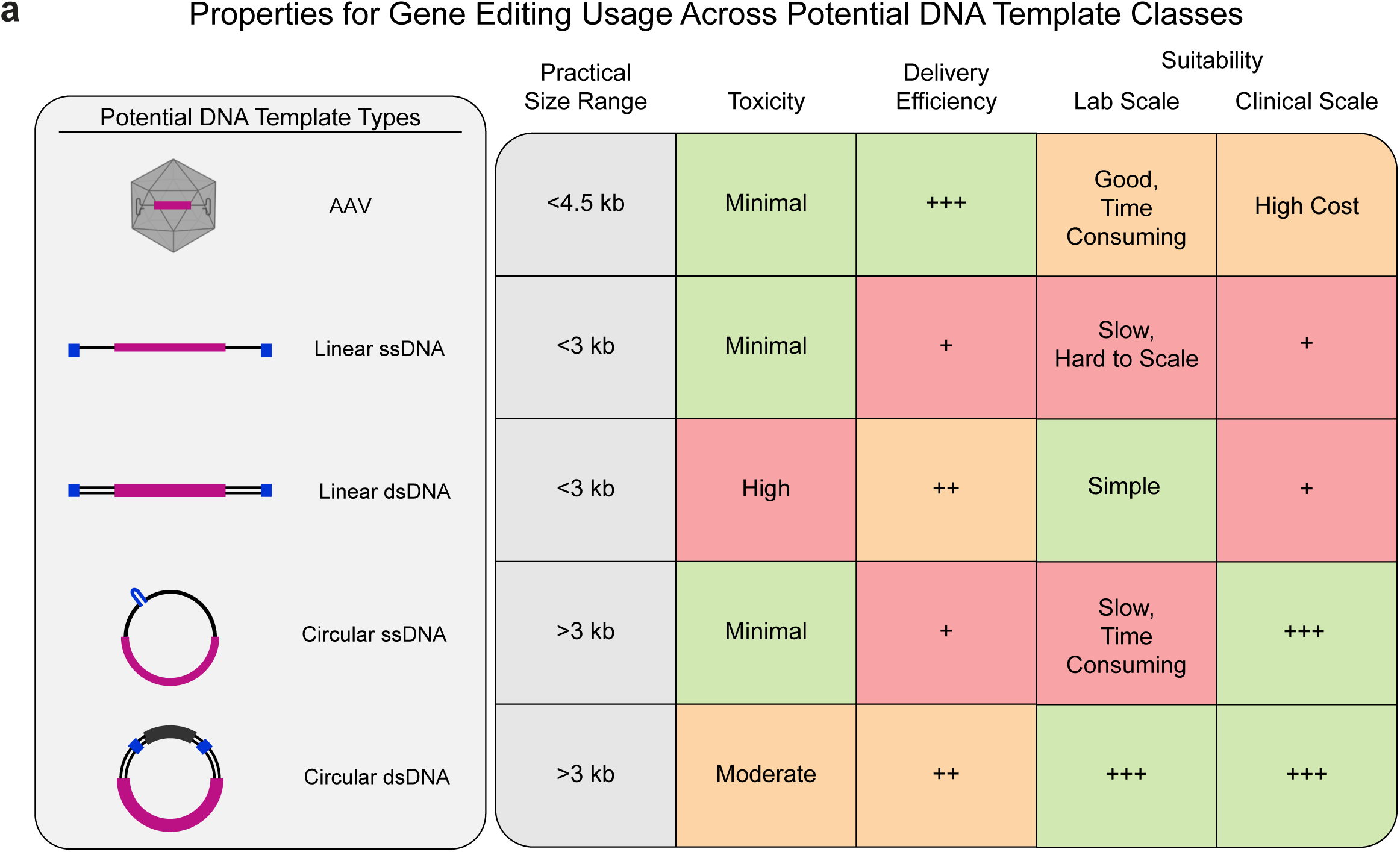
Comparative DNA Template Types for Research and Clinical Targeted DNA Integrations. **(a)** Practical size range, observed toxicity (Figure 1), delivery efficiency (**Extended Data Fig. 2**), and suitability for lab scale and clinical scale production and use (**Methods**) across a comprehensive series of potential DNA template classes, including AAVs, linear ssDNA, linear dsDNA, circular ssDNA, and circular dsDNA.

For research applications, GLIDE-editing allows for increased efficiency of DNA integrations across template sizes, with the largest increases in efficiency seen for integrations between 5kb and 10kb. The adaptability of GLIDE-editing in both primary human T cells and human iPSCs will potentially enable similar increases in editing efficiency and integration size across additional primary human cell types of interest, such as HSCs, NK cells, and B cells, as well as in human and model organism cell lines. Simple and scalable circular dsDNA plasmid templates appear to be ideal for research application of ultra-large integrations, which hopefully will broaden access to more powerful genetic manipulations. In particular, the >60% efficiency of >8kb integrations in human iPSCs without selection will enable rapid, “one-shot” production of significantly more complex DNA edits and the integration of high throughput genetic screening libraries in cells that can be differentiated into diverse downstream human cell types. GLIDE-editing thus may allow for much greater access to large scale genetic screening in both primary human cells and human iPSCs using ultra-large DNA sequences to ask more complex and powerful questions of human biologic systems.

Finally, for clinical cell therapy applications, the size of DNA sequences usable for genetically modified human cell therapies has been limited to the ∼5kb previously possible with non-viral electroporation methods, ∼4.5 kb using AAVs, or ∼5-6 kb with random integrations using lentiviral or retroviral vectors. However, increasingly sophisticated genetic circuits, with multiple coding genes and non-coding elements to encode more powerful, safer, and more controllable cellular therapies, have required larger integration sizes previously only possible at low efficiencies at research scale^24^. GLIDE-editing increases the maximum efficient size of DNA integrations in human T cells above 10 kb, effectively doubling the targetable genetic real estate available to build more effective cellular medicines. We prioritize both circular ssDNA and circular dsDNA template types as ideal for ultra-large integration methods at clinical manufacturing scale. We expect GLIDE-editing will find broad utility in enabling next-generation cell-based therapies built using ultra-large integration cassettes, as well as to increase access to scalable ultra-large gene integrations for research applications across primary human cell types and human iPSCs.

## METHODS

### Primary Human T Isolation and Culture

Human peripheral blood mononuclear cells (PBMCs) were isolated from deidentified health donor leukopacks purchased from StemCell Technologies (200-0092) or from healthy human blood donor leukoreduction system chambers (LRSCs) collected under IRB approval at the Stanford Blood Center. For LRSCs, PBMCs were isolated using density centrifugation in SepMate tubes (StemCell, 85460) with Lymphoprep (StemCell, 18061) according to manufacturer’s instructions. T cells were isolated by negative selection using EasySep Human T Cell Isolation Kit (StemCell, 17951) per manufacturer’s instructions.

With the exception of GMP-compatible scale-up experiments (described separately below), isolated CD3+ T cells were activated at 1.5e6/mL for 2 days with CTS Dynabeads CD3/CD28 (Thermo Fisher, 40203D) at a 1:1 bead to cell ratio in X-VIVO 15 medium (Lonza Bioscience, 04-418Q) supplemented with 5% fetal bovine serum (MilliporeSigma, F0926), 2 mM L-glutamine (Thermo Fisher, 2503008), and 100U/mL Penicillin-Streptomycin (Thermo Fisher, 15140122). Complete X-VIVO 15 was further supplemented with 5 ng/mL IL-7 and IL-15 (R&D Systems, 247-ILB-025 and 207-IL-050, respectively) for Fig. 1 and Extended Data Figs. 1, 2, and 4, or 50 U/mL human IL-2 (PeproTech, 200-02-1MG) for Figs. 2–4 and Extended Data Figs. 3 and 5–7. Following electroporation, T cells were split every 2-3 days using identical dilution factors (1:2 or 1:3) for all samples to prevent cell overgrowth and maintain comparable cell counts (approximately 1e6/mL).

For electroporation experiments performed on the Xenon and Maxcyte, CD3+ T cells were activated with CTS Dynabeads CD3/CD28 at a 1:1 bead to cell ratio with 20U/mL of IL-7 (Miltenyi Biotec, 170-076-114) and 100U/mL IL-15 (Miltenyi Biotec, 170-076-111) in tissue culture flasks. Following electroporation, cells were cultured at approximately 2e6 cells/mL and expanded in gas-permeable G-Rex culture vessels (Wilson Wolf, 80192M, 80240M, 80660M, RU81100) in TheraPeak X-VIVO 15 (Lonza, BP04-744Q) supplemented with 5% human AB serum (Access Cell Culture, 516-HI GI), 20U/mL of IL-7, and 100U/mL IL-15 for a 7-day expansion.

### HDRT template preparation

Long dsDNA HDRT plasmids encoding various gene inserts (**Supplementary Table 1**) and 600 bp homology arms were synthesized as gBlocks (IDT) and cloned into a pUC19 plasmid in-house or purchased directly from Genscript Biotech. All plasmids used in this study have been deposited at Addgene. These plasmids served as template for generating PCR amplicons. CTS sites were incorporated to the 5’ end of the PCR base primers. Amplicons were generated with KAPA HiFi polymerase (Kapa Biosystems, KK2602), purified by SPRI bead cleanup, and resuspended in water to 1–2 µg per µL. Amplicons were quantified using a NanoDrop spectrophotometer (Thermo Fisher, 912A1100) and purity and size were confirmed by agarose gel electrophoresis.

For ssDNA, custom ssDNA HDRTs were ordered through the Genscript service GenExact Single-Stranded DNA. To anneal complementary oligos to ssCTS templates, complementary oligos (IDT) to CTS sequences were mixed at a 4:1 molar ratio of oligos to ssDNA templates in nuclease-free duplex buffer (IDT) and were incubated at room temperature for 30 minutes. Following annealing, ssCTS templates were used immediately in electroporation experiments or aliquoted for long-term storage at −20 °C.

Custom nanoplasmids were ordered from Aldevron at 1 µg/µL via Targeted Nanoplasmid Deliverable Service. Custom circular ssDNAs were made in collaboration with Kano Therapeutics and Touchlight. Custom circular ssDNAs with CTS in hairpin loops (mbDNA) were made in collaboration with Touchlight and were manufactured using a proprietary cell-free amplification platform. For Kano circular ssDNAs templates, cssDNA design and manufacturing at scale was based on a proprietary *E. coli* cell-based fermentation process. Kano cssDNA was heated to 70 °C for 5 minutes prior to electroporation then allowed to come to room temperature. The Kano cssDNA no oligo design included single-stranded CTS reverse complements on upstream and downstream segments of the production backbone that hybridize to create a double-stranded CTS segment.

Circular dsDNA plasmid (non-nanoplasmid) templates were constructed using standard molecular cloning protocols, including PCR (NEBNext Ultra II Q5 Master Mix, New England Biolabs, M0544X), Gibson Assembly (NEBuilder HiFi DNA Assembly Master Mix, New England Biolabs, E2621L) and bacterial transformation (DH5 Alpha Competent Cells, Zymo, T3009) and purification (ZymoPURE II Plasmid Midiprep Kit, Zymo, D4201) according to manufacturers’ instructions. DNA templates were quantified using a NanoDrop spectrophotometer and diluted to 1 µg/µL for electroporation. All DNA Templates used in the study are detailed in **Supplementary Table 1**.

### Electroporation

Approximately 48 hours post-activation, T cells were mixed and separated from Dynabeads via a 2-minute incubation at room temperature using an EasySep Magnet (StemCell). Cells were counted on a Countess 3 FL Automated Cell Counter (Thermo Fisher, AMQAF1000) using Trypan Blue dye to determine viability. For each condition, 1-2e6 T cells were centrifuged at 90 × g for 10 minutes and then resuspended in 20 µL P3 Buffer (Lonza, V4SP-3960). T cells were then mixed with prepared RNP or mRNA and DNA templates.

For Fig. 1 and Extended Data Figs. 1, 2, and 4, electroporation was performed on a Lonza 4D Nucleofector Core Unit (Lonza, AAF-1003B) paired with a 96-well Unit (Lonza, AAF-1003S) using pulse code EH-115. Immediately after electroporation, 95 µL of pre-warmed complete X-VIVO 15 media was carefully added to each well and cells were recovered for 15 minutes in a 37 °C 5% CO2 incubator. T cells were then moved to an appropriate vessel to culture at 1e6 cells/mL in complete X-VIVO 15 supplemented with IL-7 and IL-15 at 5 ng/mL each.

For Figs. 2–4 and Extended Data Figs. 3, 5–7, cells were electroporated on a Lonza 4D Nucleofector Core Unit paired with a 96-well Unit using pulse code EO-151. Immediately after electroporation, 75 µL of pre-warmed complete X-VIVO 15 media was carefully added to each well and cells were recovered for 15 minutes at 37 °C in a 5% CO2 incubator. T cells were then transferred to 96-well round-bottom culture plates with 300 µL total volume of complete X-VIVO and 50U/mL IL-2.

### RNP Formulation and Electroporation

For most experiments (excluding GMP-compatible scale-up described separately below), ribonucleoproteins (RNP) were produced by either sgRNA or complexing a two-component gRNA to Cas9 with addition of ssDNAenh electroporation enhancer, as previously described^43^. Custom synthetic sgRNA from Dharmacon Horizon or Alt-R CRISPR-Cas9 sgRNA from IDT (**Supplementary Table 2**) were resuspended in 10 mM Tris-HCl (pH 7.4) with 150 mM KCl or IDT duplex buffer (IDT, 11-05-01-12) at 80 µM, and stored in aliquots at −80 °C. Synthetic CRISPR RNA (crRNA) and trans-activating crRNA (tracrRNA) were chemically synthesized (Edit-R Modified Synthetic, Dharmacon Horizon), resuspended in at a concentration of 160 µM, and stored in aliquots at −80 °C. The ssDNAenh electroporation enhancer (**Supplementary Table 2**) was synthesized by IDT, resuspended to 100 µM in water, and stored at −80 °C. To make two-component gRNA, aliquots of crRNA and tracrRNA were thawed, mixed 1:1 volume ratio, and annealed by incubation at 37 °C for 15-30 minutes to form an 80 µM gRNA solution.

To make ribonucleoprotein (RNP) (Fig. 1 and Extended Data Figs. 1, 2, and 4), sgRNA or annealed gRNA (crRNA and tracrRNA) were mixed with ssDNAenh at a 1:0.8 volume ratio prior to adding 40 µM Cas9-NLS (UC Berkeley QB3 MacroLab) at a 1:1 volume ratio to attain a molar ratio of sgRNA:Cas9 of 2:1. Final RNP mixtures were incubated at 37 °C for 15 minutes after a thorough mix. Based on a Cas9 protein, 50 pmol of RNP was used for each electroporation.

To prepare Cas9 RNPs (Figs. 2–4 and Extended Data Figs. 5–7), 0.375 µL of 200 µM tracrRNA (IDT) was mixed with 0.375 µL of 200 µM crRNA (IDT) and incubated for 15 minutes at room temperature. Next, 0.25 µL of 100 mg/mL PGA (15–50 kDa poly(L-glutamic acid), MilliporeSigma) was mixed with the complexed gRNA, followed by 0.5 µL of 40 µM Cas9-NLS, and incubated for 15 minutes at room temperature. For electroporation, µL Cas9 RNP, 2 µL of plasmid DNA template at 1 ug/µL, and 2 µL helper DNA template at 1 ug/µL were mixed with 20 µL T cells per condition.

To prepare Cas12a RNPs (Figs. 3 and 4), 0.4 µL of 200 µM Cas12a crRNA (IDT) was mixed with 0.4 µL IDT duplex buffer. Next, 0.4 µL of 100 mg/mL PGA was added and mixed by pipetting, followed by 0.8 µL of 60 µM AsUltraCas12a (UC Berkeley MacroLab), and incubated for 15 minutes at room temperature. For electroporation, 2 µL of Cas12a RNP, 2 µL of 1 ug/µL DNA template, and 2 µL of 1 ug/µL helper plasmid were mixed with 20 µL T cells per condition.

For Maxcyte and Xenon electroporation experiments (Fig. 5 and Extended Data Figs. 8 and 9), synthetic single guide RNA (sgRNA, IDT) was resuspended to 320 µM with Duplex buffer (IDT) and stored at −20 °C. For RNP formulation, sgRNA was thawed and mixed with SpyFi Cas9 nuclease (Aldevron, 9214-5MG) at a 4:1 molar ratio of sgRNA:Cas9. Final RNP mixtures were incubated at 37 °C for 15-30 minutes.

### mRNA-Based Electroporation

For mRNA electroporation with Cas9 (Fig. 1 and Extended Data Figs. 1, 2, and 4), 1 µg of Cas9 mRNA (Fig. 1F: Kano Therapeutics, Extended Data Figs 1, 2, and 4: TriLink, L-7606-100) and 1 µg of sgRNA or 1 µg of annealed tracrRNA and crRNA described above, and the indicated amount of HDRT were used per 20 µL of cells. If multiple templates were used for the same experiment, the HDRTs were normalized with duplex buffer to maintain equal volume across all conditions.

For electroporation with Cas12a (Figs. 2–4 and Extended Data Figs. 2, 5–7), editing was performed with enCas12a mRNA template, plasmid DNA template and a plasmid expressing Cas12a gRNA targeting the human *TRAC* locus (5’ GCAGACAGGGAGAAATAAGGA 3’). enCas12a mRNA was produced by *in vitro* transcription (NEB, E2080S) according to manufacturer’s instructions, with uridine replaced by N1-Methylpseudouridine-5’-Triphosphate (TriLink, N-1081-10), purified by SPRI (Cytiva, 65152105050250) cleanup, and normalized in H2O to 1 µg/µL. Prior to electroporation, 3.0 µL of enCas12a mRNA was mixed with 0.5 µL of HDR template at 1 µg/µL, 0.5 µL of gRNA expression plasmid at 1 µg/µL, and 1.5 µL of helper plasmid at 1 µg/ µL, and subsequently combined with T cells in P3 Buffer.

### Clinical-Scale Electroporation

For Xenon and Maxcyte experiments (Fig. 5 and Extended Data Figs. 8 and 9), approximately 48 hours post-activation, T cells were mixed and separated from Dynabeads via a 2-minute incubation at room temperature using an EasySep Magnet (StemCell).. Cells were left to rest in a tissue culture flask at 37 °C for 1 hour prior to electroporation. After resting, the cells were centrifuged for 10 minutes at 90 × g and resuspended with CTS Xenon Genome Editing Buffer (Gibco, A4998001) or Maxcyte Electroporation Buffer (Maxcyte, EPB-1). HDRT and RNP were mixed and incubated for at least 10 minutes before being combined with cells.

For electroporation on Maxcyte, 2-5e6 cells were electroporated with 2 μM of RNP and the indicated amount of HDRT using R50×8 process assembly on Maxcyte GTx using pulse code ETC 4-2. Immediately post-electroporation, 20 µL of prewarmed cytokine-free complete TheraPeak X-VIVO 15 was added to each well without disturbing the cells. Cells were then incubated for 30 minutes at 37 °C before being transferred to G-Rex culture vessels and cultured at 2e6 cells/mL in complete TheraPeak X-VIVO 15 with 20 IU/mL IL-7 and 100 IU/mL IL-15.

For electroporation on Xenon, 50e6 cells were electroporated with 1 µM of RNP and an indicated amount of HDRT in a final volume of 1 mL in a CTS Xenon Single Shot Electroporation Chamber (Thermo Fisher, A50305). Following electroporation, cells were left to rest for 1-2 minutes in the cuvette before being transferred to a tube containing 1 mL of prewarmed cytokine-free complete TheraPeak X-VIVO 15. The cells were incubated for 30 minutes at 37 °C before being transferred to G-Rex culture vessels and cultured at 2e6 cells/mL in complete TheraPeak X-VIVO 15 with 20 IU/mL IL-7 and 100 IU/mL IL-15.

### Human iPSC Culture and Electroporation

Human iPSCs were purchased from commercial vendors: Fibroblast-derived KOLF2.1J iPSCs from The Jackson Laboratory (JIPSC001000) and blood-derived SCTi004 iPSCs from StemCell Technologies (200-0769). Cells were maintained on 5 µg/mL CellAdhere Laminin-521 coated plates following manufacturer’s protocol (StemCell Technologies, 200-0117) in complete eTeSR medium (StemCell Technologies, 100-1215). iPSCs were passaged every 4-5 days upon reaching 70-90% confluency by dissociation using TrypLE Express (Thermo Fisher, 12-604-021) for 5 minutes in a 37 °C 5% CO2 incubator and were seeded for the first 24 hours in eTeSR supplemented with 1 µM H 1152 dihydrochloride rho-kinase inhibitor (Tocris, 24-141) to enhance cell survival of single cells. Subsequent days, medium was exchanged daily with eTeSR without H 1152.

For electroporation experiments, cells were dissociated and counted on a Countess 3 FL Automated Cell Counter using Trypan Blue dye 1e6 viable iPSCs per editing condition were centrifuged at 90 × g for 5 minutes and resuspended in 20 µL P3 Buffer. iPSCs were then combined with the prepared mRNA+DNA mixture: 3 µL of 1 µg/µL Cas9 mRNA, 0.5 µL of 1 µg/µL HDR template, 0.5 µL of 1 µg/µL gRNA expression plasmid, and 1.5 µL of 1 µg/µL helper plasmid. Cas9 mRNA was produced by *in vitro* transcription (NEB, E2080S) according to manufacturer’s instructions, with uridine replaced by N1-Methylpseudouridine-5’-Triphosphate (TriLink, N-1081-10), purified by SPRI cleanup, and normalized in H2O to 1 µg/µL. Electroporation was performed on a Lonza 4D Nucleofector Core Unit paired with a 96-well Unit using pulse code EO-151. Immediately after electroporation, 75 µL of room temperature eTeSR + H 1152 was carefully added to each well and cells were recovered for 15 min in a 37 °C 5% CO2 incubator. iPSCs were then transferred to a Laminin-521 coated 6 well plate in 2 mL eTeSR + 1 µM H 1152 per well. The following day, medium was exchanged with eTeSR without H 1152, and medium was changed daily until cells were ready for analysis. On Day 6 post-electroporation, cells were dissociated and stained with an APC anti-human CD271(NGFR) antibody (BioLegend, 345108, 1:100 dilution) and analyzed by flow cytometry using a ZE5 Cell Analyzer (Bio-Rad).

### Flow cytometry

Flow cytometry analysis was performed on an Attune NxT flow cytometer with a 96-well autosampler (Thermo Fisher Scientific) or ZE5 Cell Analyzer (Bio-Rad). Unless otherwise indicated, cells were collected 5-7 days post-electroporation, resuspended in FACS buffer (PBS (Corning, 21040CM), 2% fetal bovine serum (MilliporeSigma, F0926), and 0.2% EDTA (Invitrogen, 15575020)) and stained with Ghost Dye Red 780 (Tonbo, 13-0865-T100) and the indicated cell-surface markers (**Supplementary Table 3**). All samples were washed with FACS buffer prior to staining. To stain for BCMA-CAR, cells were incubated with 0.3 µg BCMA recombinant protein conjugated to biotin (Biotinylated Human BCMA / TNFRSF17 Protein, His, Avitag, AcroBioSystems, BCA-H82E4_200ug) for 15 minutes at room temperature prior to surface cell staining. Knockout was assessed using anti-TCRα/β antibody while knock-in was assessed using anti-Streptavidin. To stain for PDAC-synNotch, cells were incubated with 50x diluted Fc block (BioLegend, 422302) in FACS buffer at room temperature for 10 minutes prior to surface staining. Knock-in was assessed with anti-Whitlow and anti-G4S was used to assess CAR leakage. RQR8 or anti-CD34 was used to detect CD19-CAR and anti-CD3 was used to detect knockout. For surface marker staining, all samples were incubated with antibodies diluted in FACS buffer for 20 minutes to 1 hour at 4 °C followed by two washes. To obtain comparable live cell counts between conditions, events were recorded from an equivalent fixed volume for all samples. Analysis was done using FlowJo v10 software. All gating strategies included exclusion of subcellular debris, singlet gating, and viable cells. All antibodies used in the study are described **in Supplementary Table 3**.

### AAV Template Production

AAV-ITR plasmids containing the BCMA-CAR and TRAC-targeting homology arms for homology directed repair were used as previously described^39^. The AAV-ITR-containing plasmid was packaged into AAV6 using polyethylenimine-based co-transfection of HEK293T cells with pHelper and pAAV Rep-Cap plasmids. Viral particles were extracted from cells and purified using iodixanol-based density gradient ultracentrifugation. AAV titration was performed by qPCR after treating samples with DNase I (MilliporeSigma, 4716728001) and Proteinase K (Roche, 3115887001), using primers targeting the left homology arm (Forward: CTTTGCTGGGCCTTTTTCCC, Reverse: CCTGCCACTCAAGGAAACCT). qPCR was performed using SsoFast Eva Green Supermix (Bio-Rad, 1725201) on a StepOnePlus Real-Time PCR System (Applied Biosystems).

Primary human T cells were isolated, activated, and electroporated with preassembled Cas9 RNPs as described above. Following electroporation, cells were rescued with 90 µL prewarmed complete X-VIVO 15 without added serum and incubated for at least 15 minutes at 37 °C. Recombinant AAV6 donor vector was added to the culture 30–60 min after electroporation at the multiplicity of infection specified, and cells were incubated overnight in serum-free X-VIVO 15 with IL-7 and IL-15 at 5 ng/mL. The next day, the serum-free AAV-containing medium was removed, and cells were resuspended in fresh complete X-VIVO 15 with IL-7 and IL-15 at 5 ng/mL and expanded using standard culture conditions (37 °C, 5% CO2, and complete medium replenished as needed to maintain a density of 1e6 cells/ mL every two to three days). Knockout and knock-in efficiency were evaluated by staining for the TCR with an anti-TCRα/β antibody (Miltenyi Biotec) and staining for the CAR with recombinant BCMA protein (Biotinylated Human BCMA / TNFRSF17 Protein, His, Avitag, AcroBioSystems BCA-H82E4_200ug). Flow cytometry was conducted on a Attune NxT flow cytometer with a 96-well autosampler (Thermo Fisher).

### DNA Template Delivery and Half-life Quantification

At the indicated timepoints, for each treated sample and no treatment control, 1.5e6 cells were centrifuged at 450 × g for 5 minutes, washed with 1x PBS, and stored at −80 °C. To extract DNA, cell pellets were thawed on ice and extracted in accordance with manufacturer’s protocol using QIAamp DNA Mini Kit (Qiagen, 51306). Following extraction, samples were eluted in nuclease-free water (Invitrogen, 10977-015). Samples were then quantified using the Qubit dsDNA Broad Range Kit (Thermo Fisher, Q32850) and normalized to 5 ng/µL. qPCR was used to quantify the amount of template per sample. Custom BCMA probe and primers developed by Thermo Fisher were used at final concentrations of 250 nM and 900 nM respectively (**Supplementary Table 4**). ActB was used as the housekeeping gene and measured using ActB TaqMan Gene Expression Assay (Thermo Fisher, 4351370). Primer and probes were diluted using nuclease-free water and TaqMan Fast Advanced Master Mix (Thermo Fisher, 4444963). Wild-type DNA for each donor was run to generate ActB standard curves. 6-step 10x serial dilutions of each HDRT template were made starting with 0.1 pmol/µL and run with BCMA primer and probes. 2 µL of sample and 8 µL of primer probe master mix were loaded into 384 well plates (Thermo Fisher, 164610). Standards and samples were run in triplicate. Before running, plates were centrifuged for 1 minute at 3000 RPM. Plates were run on QuantStudio 5 (Thermo Fisher, A34322). The amount of template was calculated using the corresponding template standard curve and then normalized by the number of genomes calculated based on ActB Ct, ActB standard curves, and mass of the human genome.

## SUPPLEMENTARY INFORMATION

Supplementary Table 1. DNA Template Sequences used in study.

Supplementary Table 2. gRNA and Electroporation enhancer sequences.

Supplementary Table 3. Antibodies used in study.

Supplementary Table 4. DNA primers used in study.

## DATA AVAILABILITY

All underlying data presented are available upon request to corresponding authors.

## ACKNOWLEDGEMENTS

We thank the members of the Allen, Shy, and Roth Labs for stimulating discussions. The Roth Lab has received sponsored research support from an NIH Director’s New Innovator Award (DP2 CA311217), the NCI (K08 CA286740), a Burroughs Wellcome Career Award for Medical Scientists, a Senior Fellow award from the Parker Institute of Cancer Immunotherapy, an Innovation Investigator Award from the Arc Institute, a The Cancer League Research Grant Program Award, a Gates Foundation Innovation Pilot Award, and received support from the Weill Foundation and Northpond Ventures. T.L.R. is a Member of the Parker Institute of Cancer Immunotherapy, an Affiliate Investigator of the Arc Institute, and an Investigator at the Weill Cancer Hub West. L.C., M.A.G., W.A.L., R.A., G.A. and B.R.S. are supported by the CRISPR Cures for Cancer initiative by UCSF, Gladstone Institutes and Innovative Genomics Institute at UC Berkeley. B.R.S is supported by NIH grants K08CA273529 and L30TR002983, California Institute of Regenerative Medicine (CIRM) grants INFR5-14719 and CLIN1-15060, the UCSF CRISPR Cures for Cancer Initiative, the Stephen and Nancy Grand Multiple Myeloma Translational Initiative, and the UCSF Living Therapeutics Initiative. G.M.A is supported by NIH grant K08CA259610, California Institute of Regenerative Medicine (CIRM) grant TRAN1-16998, the Mark Foundation, the Lydia Shorenstein Donor Advised Fund, the UCSF CRISPR Cures for Cancer Initiative, and the UCSF Living Therapeutics Initiative.

## CONTRIBUTIONS

Research design, C.K., L.C., M.G.A.., K.L, T.T., M.F., R.A., G.M.A., B.R.S., and T.L.R. Writing, C.K., L.C., R.A., G.M.A., B.R.S., and T.L.R. with input from all authors. Lonza T cell knock-in experiments and flow cytometry readouts, C.K., L.C., M.G.A.. iPSC electroporations, C.K. T.L.R. GMP-compatible manufacturing experiments, K.L., T.T. and J.L. Helper plasmid optimization experiments, C.K. and T.L.R. Large knock-in sequence optimization and GLIDE editing protocol, C.K. and T.L.R. In vitro cytotoxicity assays, M.F. and M.G.A. In vivo experiments, M.F. In vitro induction experiments, M.G.A. DOE experiments, K.L., T.T., Y.J., N.P., N.A., N.F., M.D., AAV production, L.C. and L.W. DNA stability assay. Supervision, O.K., R.A., G.M.A, T.R., B.R.S. cssDNA design and production, Kano Therapeutics Inc. and Touchlight.

## EXTENDED DATA FIGURE LEGENDS

**Extended Data Figure 1.**
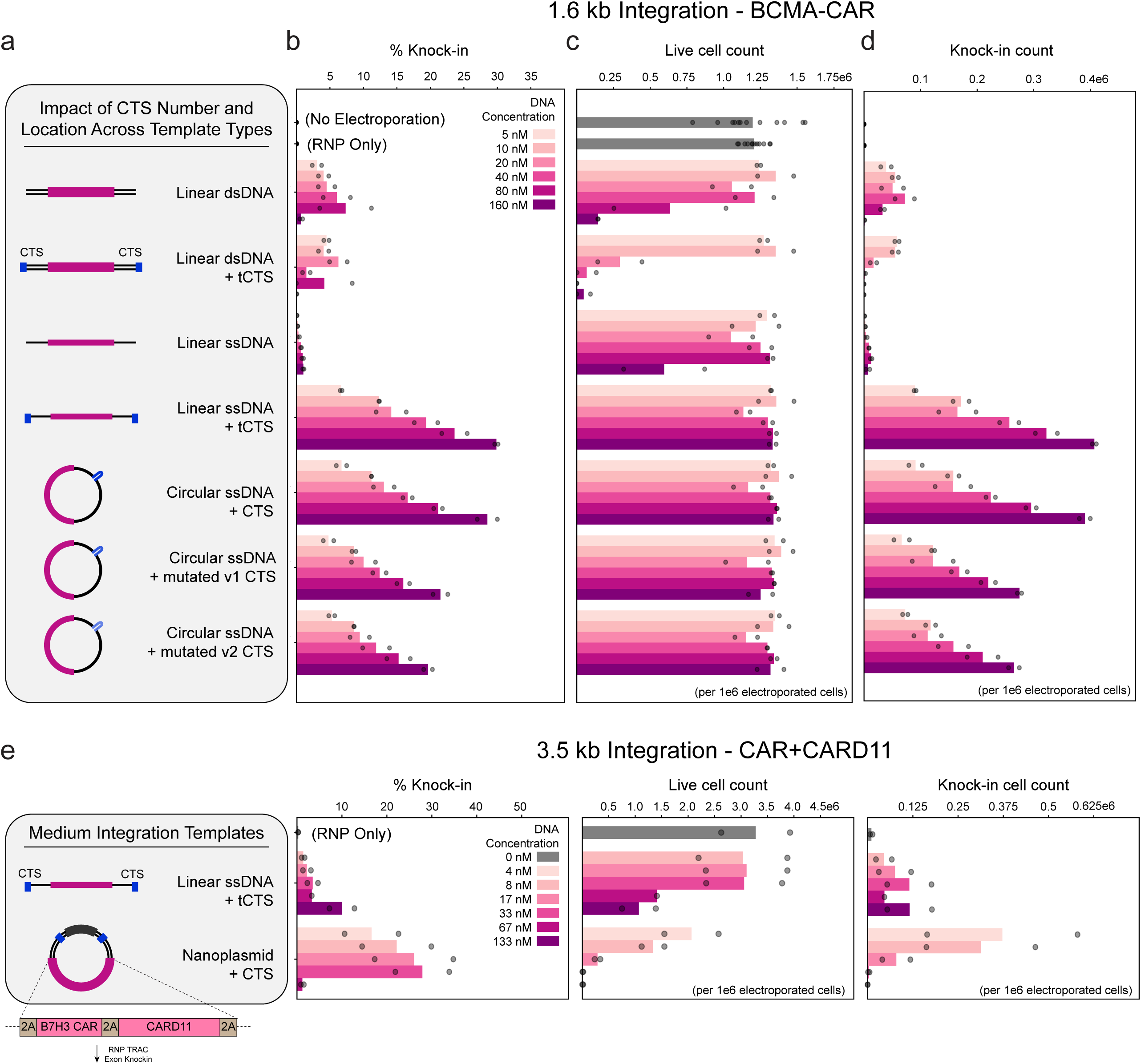
Optimizing CTS engineering and electroporation workflow. **(a)** Knock-in strategy and designs for BCMA-CAR (1.6 kb integration) at concentrations 5nM-160nM with corresponding **(b)** knock-in efficiency, **(c)** live cell count, and **(d)** knock-in cell count 7 days post electroporation for the following templates respectively: dsDNA, dsDNA + tCTS, ssDNA, ssDNA + tCTS, circular ssDNA + CTS, circular ssDNA + mutated v1 CTS v1, circular ssDNA + mutated v2 CTS. Circular cssDNA was produced and provided by Touchlight. **(e)** Knock-in strategy and designs for CAR-CARD-11 (3.5 kb integration) using linear ssDNA + tCTS template or a nanoplasmid + CTS template at concentrations 5nM-160nM with corresponding knock-in efficiency, live cell count, and knock-in cell count 7 days post electroporation.

**Extended Data Figure 2.**
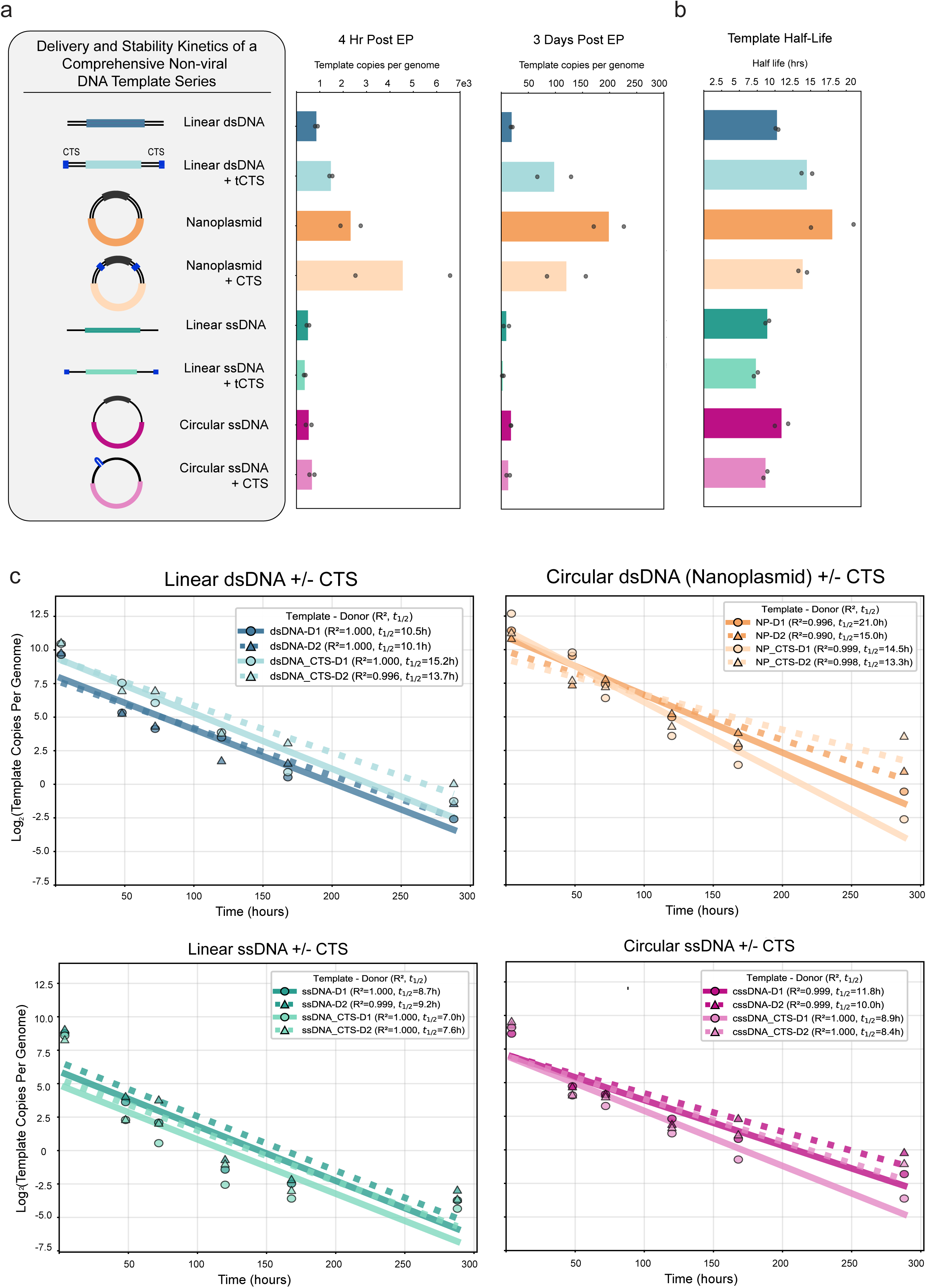
Efficiency of intracellular delivery across template types. **(a)** Designs for BCMA-CAR (1.6 kb integration) HDRTs variants tested and corresponding template copies per genome measured at 4 hours and 3 days post electroporation **(b)** and half-life measured by qPCR. **(c)** Template copies per genome measured over time at 4 hours, 2 days, 3 days, 5 days, 7 days, and 12 days with R2 and half-life values corresponding to each donor. Each experiment was performed with T cells from two independent healthy human blood donors represented by individual dots plus mean. tCTS, truncated Cas9 target site. CTS, Cas9 target site. NP, nanoplasmid. cssDNA, circular cssDNA.

**Extended Data Figure 3.**
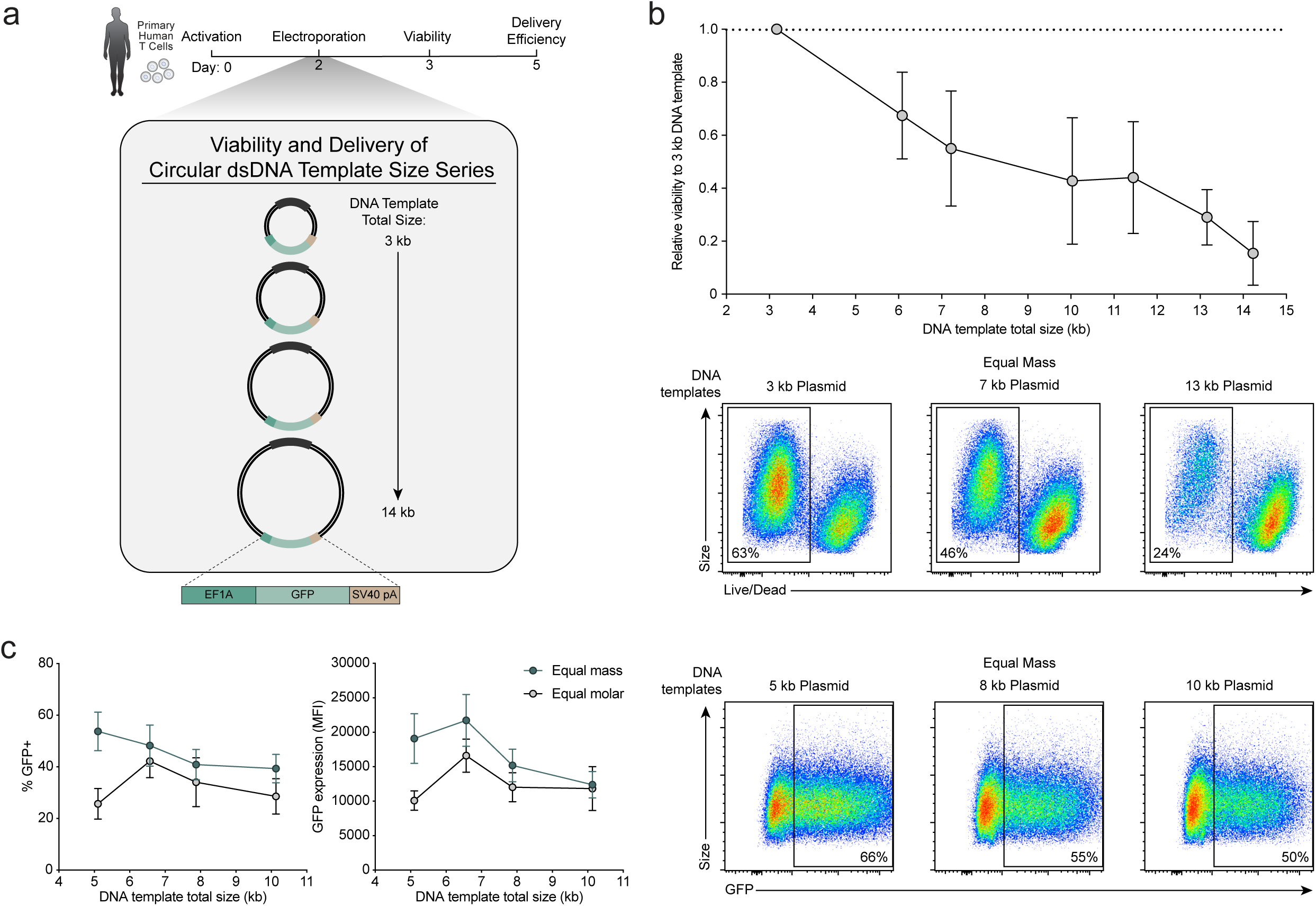
Maximal size of circular dsDNA delivery into human T cells. **(a)** Timeline of electroporation of primary human T cells with episomal GFP expression plasmids of increasing size (circular dsDNA templates). **(b)** Live cell counts 24 hours post-electroporation relative to small DNA template (3 kb total template) (*n* = 3-5 donors, mean ± s.d.). Representative flow plots showing decreased viability as template size increases. **(c)** DNA template delivery, with plasmid size series delivered with equal mass or molar. Delivery efficiency and MFI in GFP^+^ cells 5 days post-electroporation (*n* = 3 donors, mean ± s.d.). Representative flow plots showing decreased delivery as template size increases.

**Extended Figure 4.**
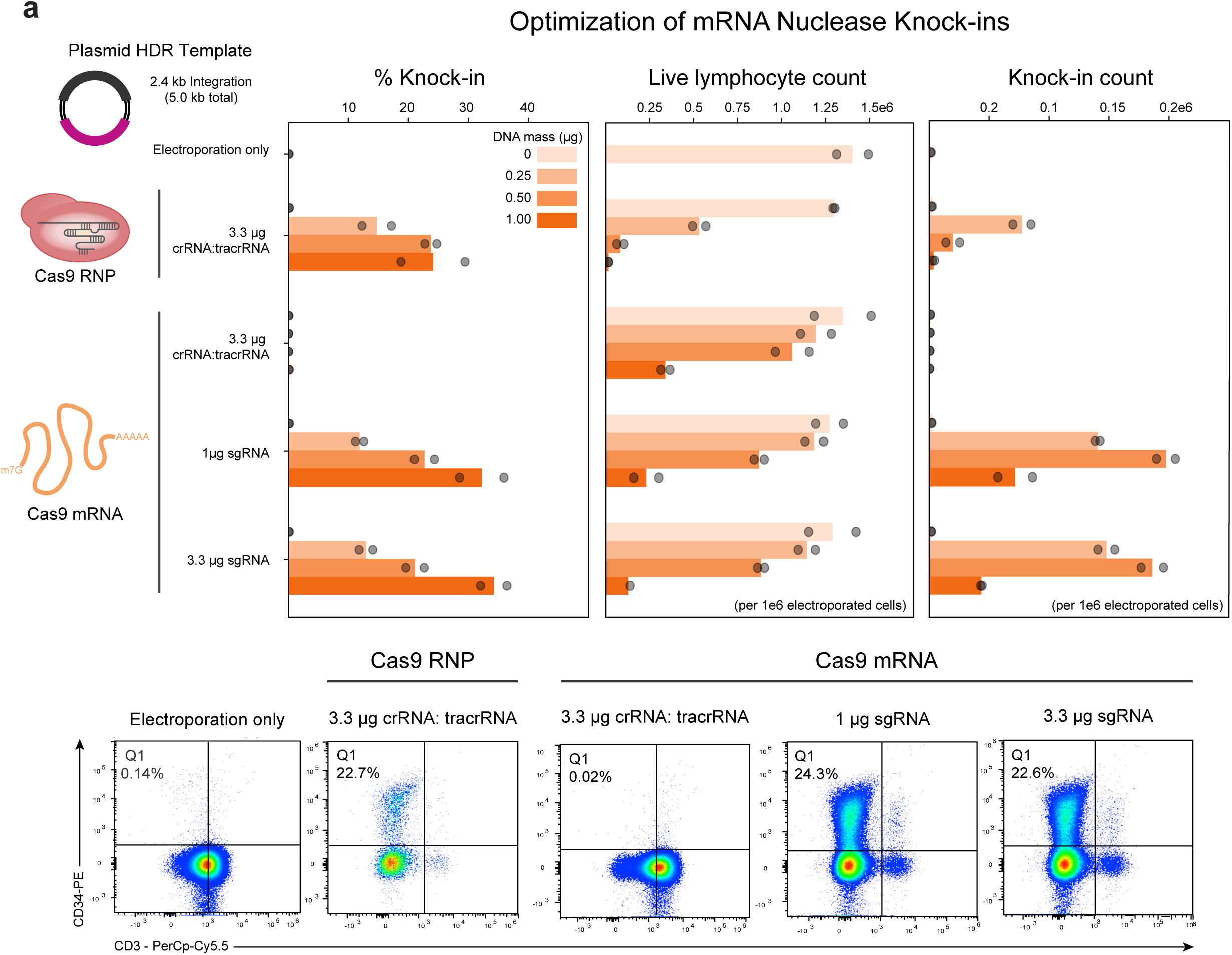
mRNA enhances large dsDNA delivery and editing efficiency. **(a)** Comparison of Cas9 RNP and Cas9 mRNA with either crRNA: tracrRNA or sgRNA for a circular dsDNA plasmid HDR template encoding a CD19-28Z CAR (2.4 kb integration, 5.0 kb total template size) at 0-1.00 µg with corresponding knock-in efficiency, live cell count, knock-in cell count, and representative flow plots for the 0.5 ug condition at day 7 post activation, equivalent to 5 days post electroporation. Each experiment was performed with T cells from two independent healthy human blood donors represented by individual dots plus mean. CTS, Cas9 target site. RNP, ribonucleoprotein. crRNA, CRISPR RNA. tracrRNA, trans-activating RNA. sgRNA, single-guide RNA.

**Extended Data Figure 5.**
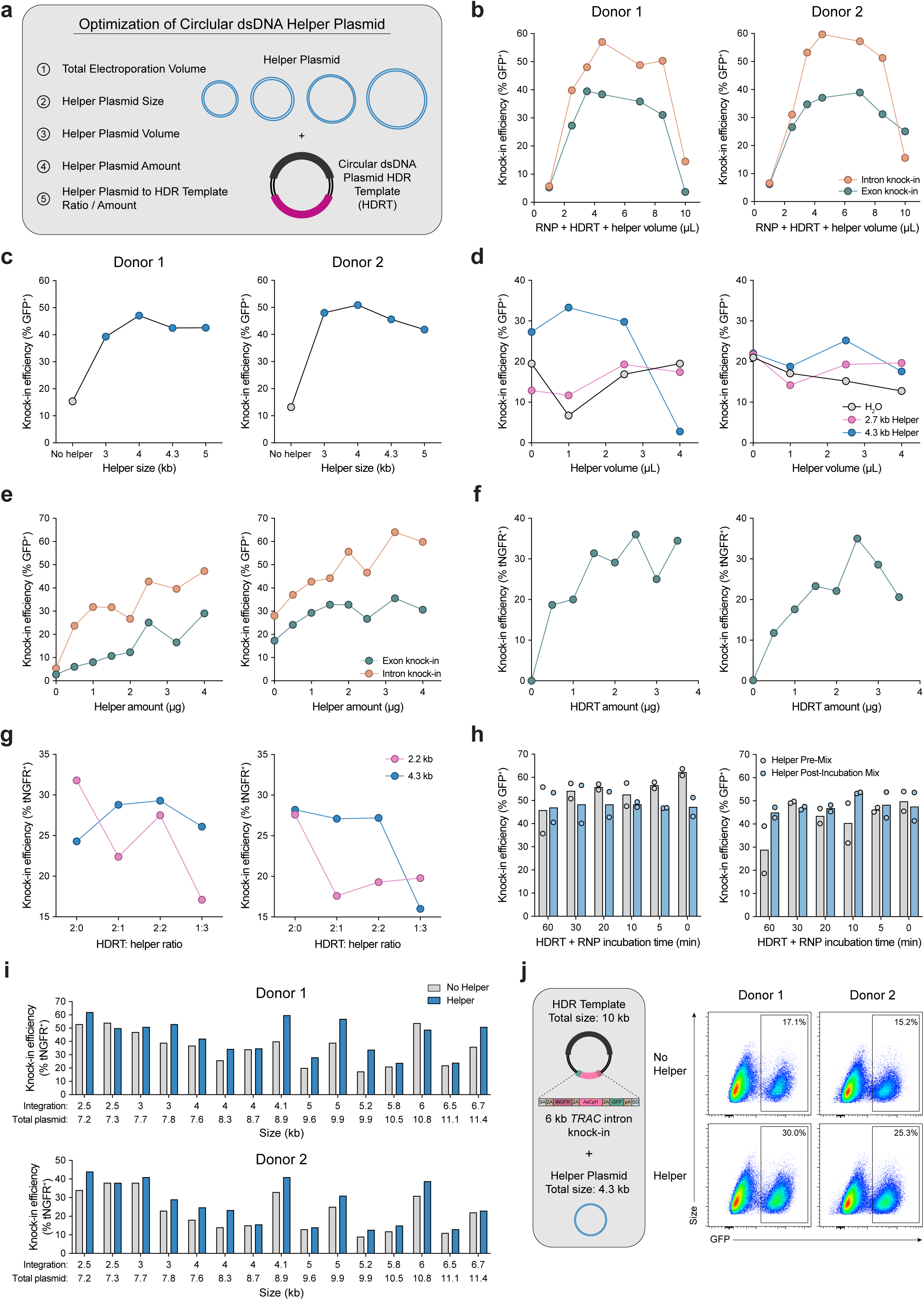
Optimization of small helper plasmid assisted large DNA template delivery and integration. **(a)** Optimization of circular dsDNA helper plasmid to enhance delivery and knock-in efficiency. **(b)** Optimization of combined homology-directed repair template (HDRT), helper plasmid and RNP volume. Knock-in efficiency (4.1 kb and 6.0 kb integrations) at TRAC intron or exon sites. **(c)** Optimization of helper plasmid size. Knock-in efficiency (5.2 kb integration) with co-delivery of helper plasmids with varying sizes. **(d)** Optimization of helper plasmid volume (2.7 or 4.3 kb total template) for efficient knock-in (6 kb integration). **(e,f)** Optimization of helper plasmid amount (4 kb total template) **(e)** and HDRT amount **(f)** for maximal knock-in efficiency (5.2 kb integration) at TRAC intron or exon sites. **(g)** Effect of HDRT-to-helper plasmid ratio on knock-in efficiency (4.1 kb integration). **(h)** Effect of mixed HDRT and RNP incubation time, with helper plasmid added before or after incubation, on knock-in efficiency (5.2 kb integration). **(i)** Knock-in efficiency across HDRT sizes with co-delivery of helper plasmid (4.3 kb total template) compared to template-only controls. **(j)** Representative flow cytometry plots showing knock-in efficiency with or without helper plasmid (4.3 kb total template). **(b-j)** Knock-in efficiency measured in edited (GFP^+^ or tNGFR^+^) cells 5 days post-electroporation in two donors.

**Extended Data Figure 6.**
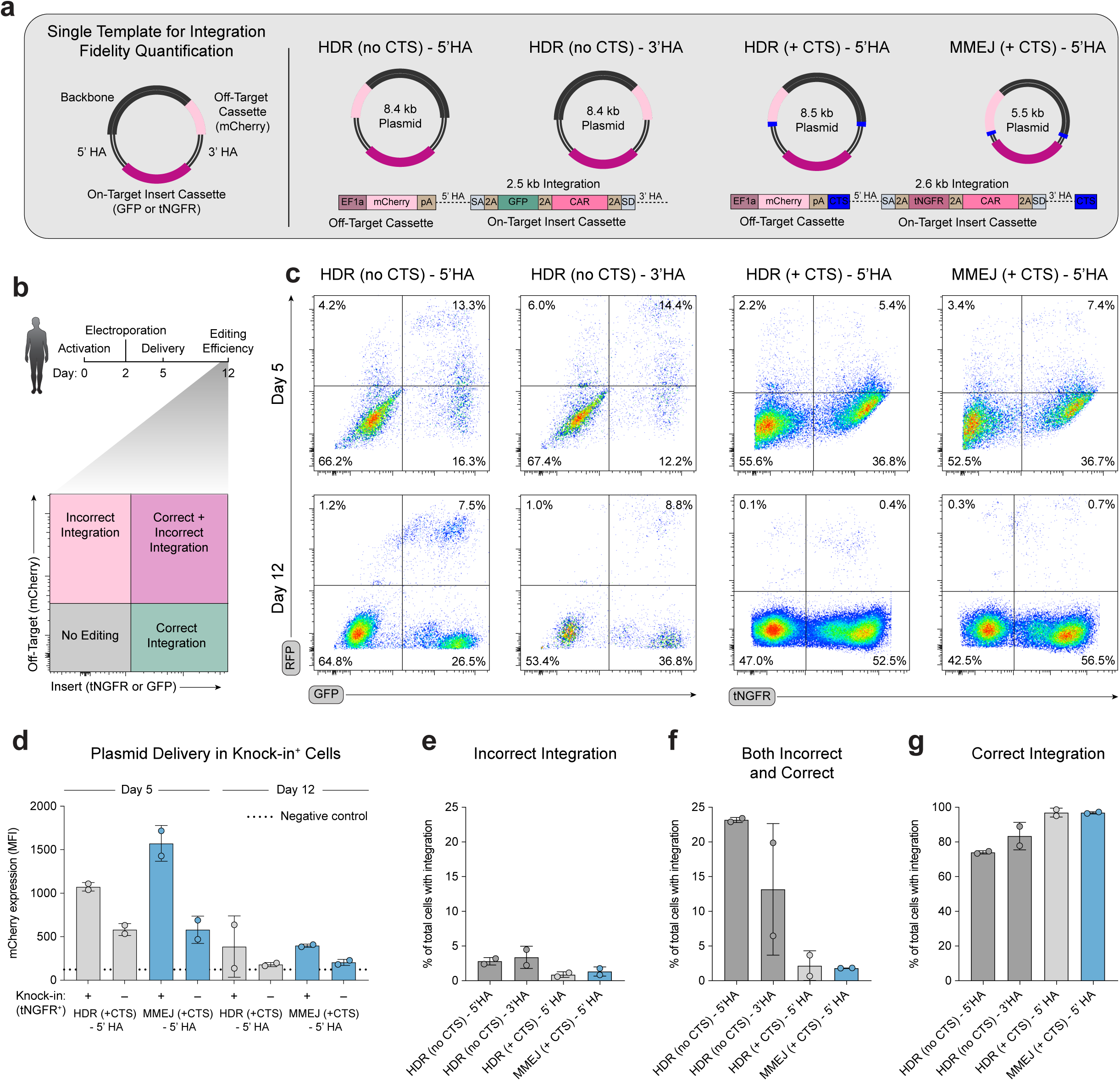
Single cell quantification of off-target integrations of large templates. **(a)** Plasmid series containing an off-target integration control cassette outside of homology arms of a multicistronic cassette integrated at the TRAC locus. **(b)** Experimental timeline with delivery measured by episomal mCherry expression. Integration measured in GFP^+^ or tNGFR^+^ cells with correct integration and mCherry^+^ cells with incorrect integration of the off-target cassette. **(c)** Representative flow cytometry plots of episomal mCherry washout between day 5 and 12. **(d)** DNA template delivery on day 5 and washout by day 12. mCherry MFI measured in edited (tNGFR^+^) or unedited (tNGFR^-^) cells and compared to unedited (tNGFR^-^) cells from a negative control (HDR template, no off-target cassette). **(e)** Incorrect integration, quantified as the proportion of mCherry^+^ cells among total cells with integration. **(f)** Concurrent incorrect and correct integration, quantified as the proportion of mCherry^+^ tNGFR^+^ or mCherry^+^ GFP^+^ cells among total cells with integration. **(g)** Correct integration, quantified as the proportion of tNGFR^+^ or GFP^+^ cells among total cells with integration. **(e-g)** Integration measured by flow cytometry 10 days post-electroporation (*n* = 2 donors, 2 technical replicates, mean ± s.d.).

**Extended Data Figure 7.**
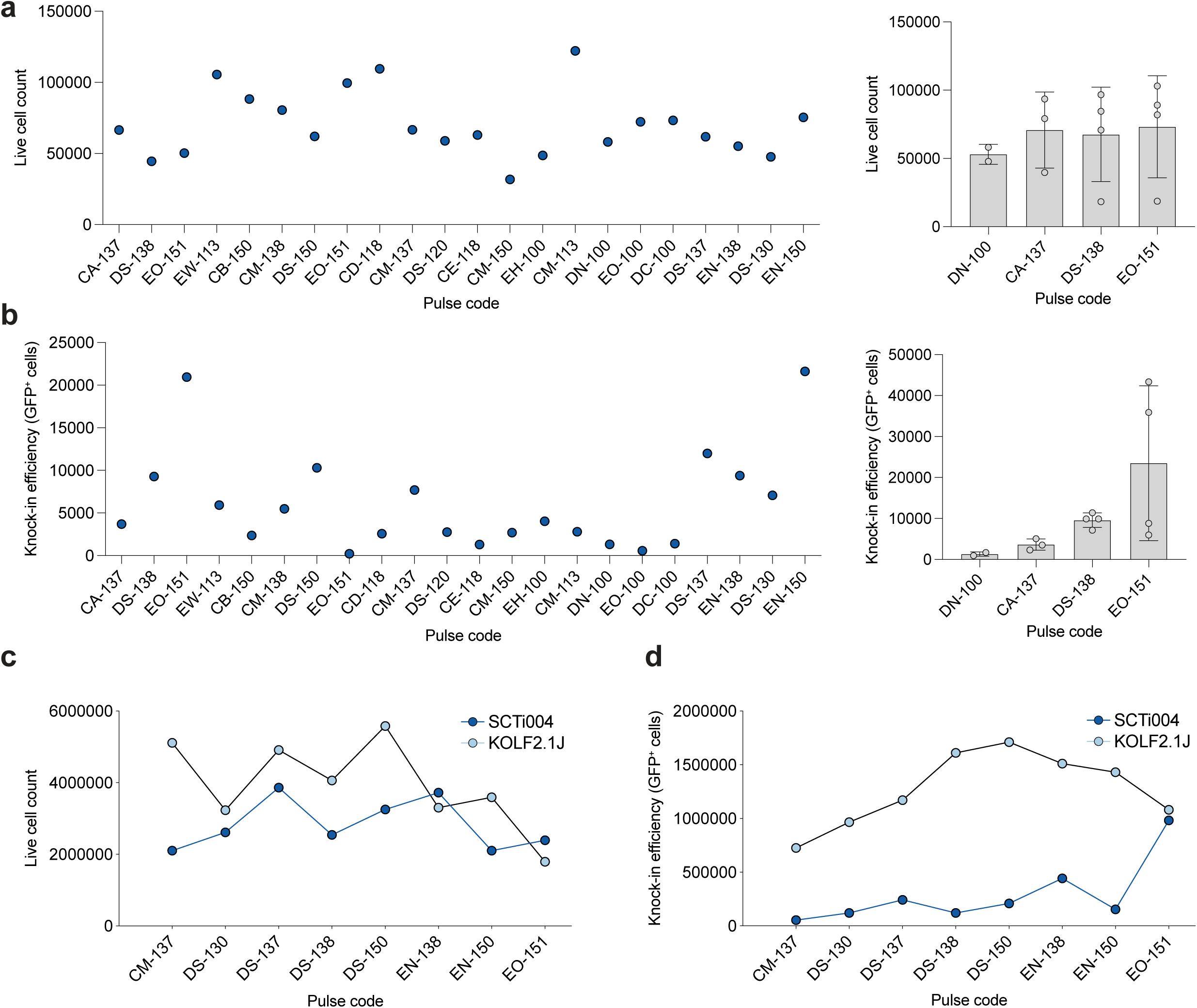
Optimization of ultra-large DNA integrations in human iPSCs. **(a)** Live cell counts and **(b)** knock-in efficiency with a series of electroporation pulse codes in iPSCs (SCTi004). Live and edited (GFP^+^) cells measured 6 days post-electroporation (2-4 technical replicates for top pulse codes, mean ± s.d.). **(c)** Live cell count and **(d)** knock-in efficiency with a series of top electroporation pulse codes in two iPSC lines (SCTi004 and KOLF2.1J). Live and edited cells (GFP^+^) measured 6 days post-electroporation.

**Extended Data Figure 8.**
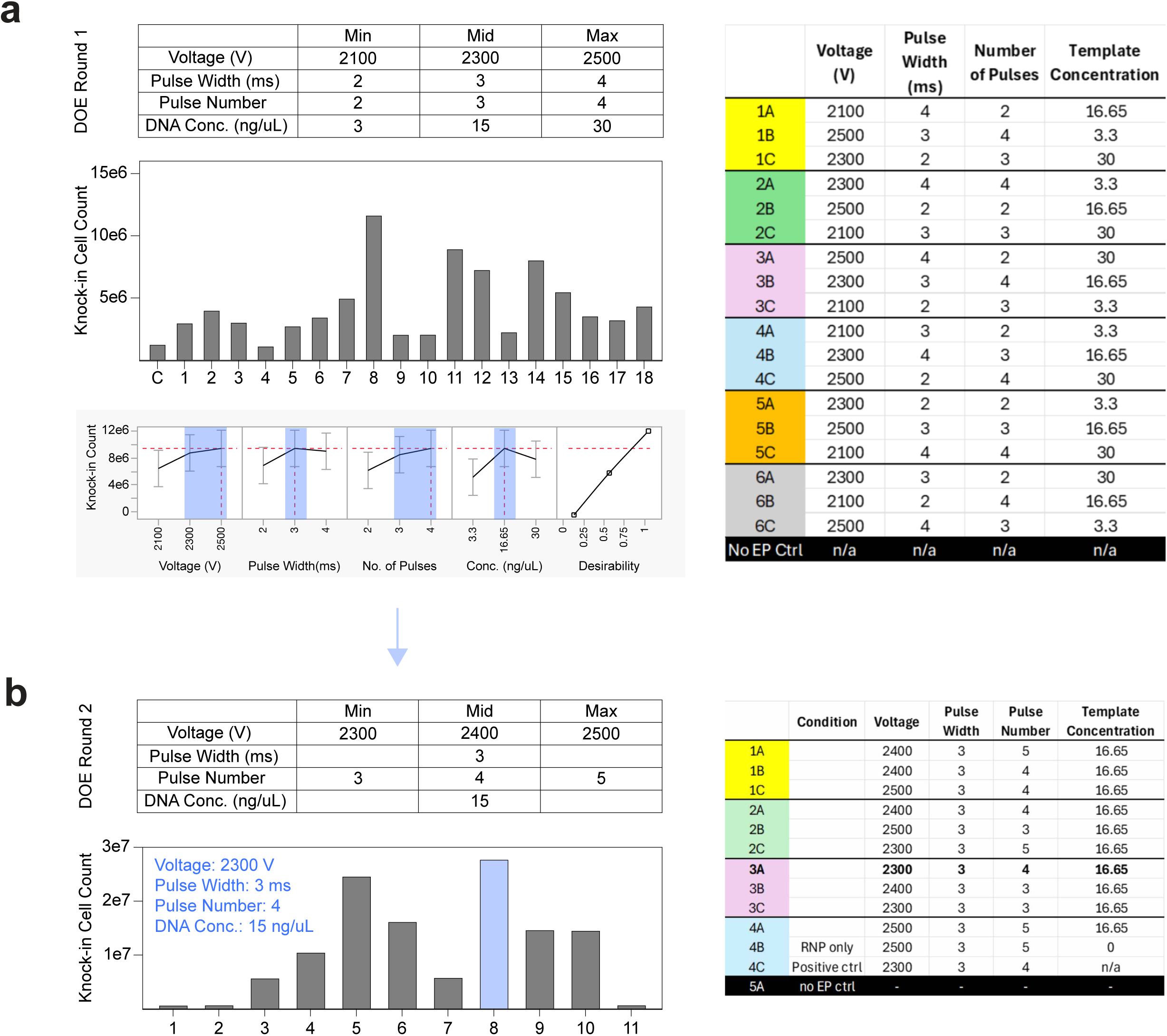
Parameter optimization of clinical manufacturing of ultra-large integrations in human T cells. **(a)** Design-of-experiments (DoE) optimization Round 1. A definitive screening design was performed on the Xenon clinical-scale electroporation system to evaluate the effects of voltage (2100-2500 V), pulse width (2-4 ms), pulse number (2-4), and circular dsDNA nanoplasmid template concentration (3-30 ng/µL) on knock-in cell yield. Nanoplasmid DNA template encoded a synNotch circuit targeted to the *TRAC* locus (5.6 kb integration, 7.3 kb total template size). Top, parameter ranges and experimental matrix tested. Middle, total viable knock-in cell counts for each condition (C, control). Bottom, main effects analysis showing the relative contribution of each parameter to knock-in yield and an overall desirability function. JMP prediction profiler predicted increased voltage and template concentration improved knock-in cell recovery within the tested range, while intermediate pulse width and pulse number conditions balanced efficiency and viability. These data defined a preliminary optimal region for further refinement. **(b)** DoE Round 2. Focused optimization using narrowed parameter ranges centered on Round 1 optima (voltage 2300-2500 V; pulse width 3 ms; pulse number 3-5; template concentration 15 ng/µL). Left, updated parameter space and resulting viable knock-in cell counts across conditions. The optimal condition (highlighted) was 2300 V, 3 ms pulse width, 4 pulses, and 15 ng/µL template DNA, yielding the highest recovery of viable knock-in cells. Right, experimental design matrix including RNP-only and no-electroporation controls.

**Extended Data Figure 9.**
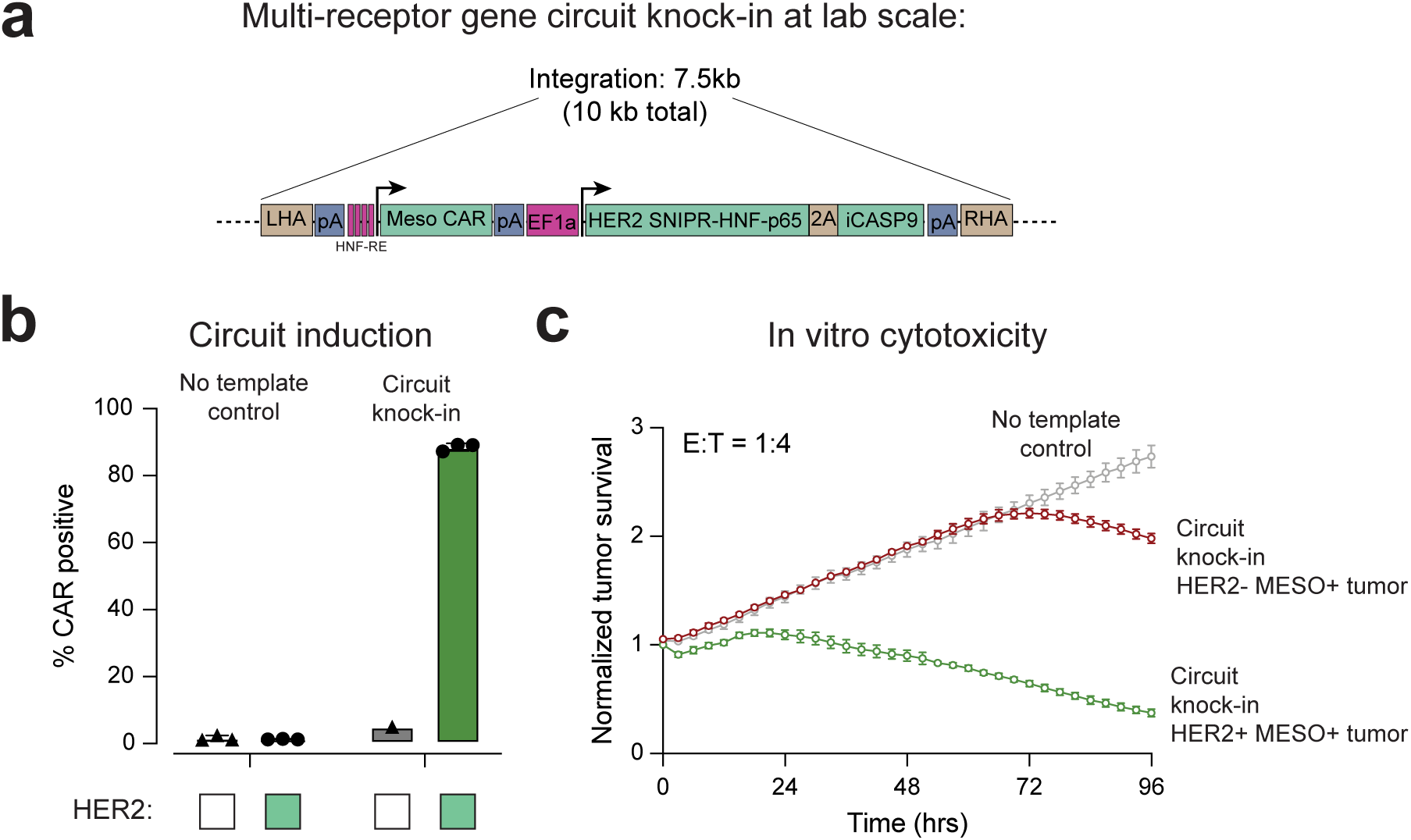
Functional validation of lab-scale ultra-large integrations in human T cells. **(a)** Non-viral integration of a logic-gated circuit (7.5 kb integration) at the *TRAC* locus using dsDNA plasmid (10 kb total size) into primary human T cells. The circuit contains a constitutively active synthetic intramembrane proteolysis receptor (SNIPR) and inducible anti-mesothelin CAR. **(b)** In vitro CAR induction assay. 1e5 circuit engineered or no template control T cells were co-cultured with 1e5 K562s with or without HER2 antigen expression for 48 hours. CAR expression was measured by flow cytometry showing specific induction of CAR expression in circuit T cells in presence of HER2+ target cells. Mean +/- s.d. depicted, n=3. **(c)** In vitro T cell cytotoxicity assay. 5e3 circuit T cells were co-cultured with 2e4 GFP+ AsPC-1 tumor cells (E:T = 1:4) on an Incucyte Live Cell Imaging system. Survival of GFP+ tumor cells was measured directly, showing specific lysis of HER2+ AsPC-1 tumor cells and not HER2-AsPC-1 tumor cells. No template knock-in T Cells were used as a control. Mean +/- s.d. of GFP signal normalized to time 0 is depicted, n=3.

## REFERENCES

1. Naldini, L. et al. In vivo gene delivery and stable transduction of nondividing cells by a lentiviral vector. Science 272, 263–267 (1996).

2. Sadelain, M., Wang, C. H., Antoniou, M., Grosveld, F. & Mulligan, R. C. Generation of a high-titer retroviral vector capable of expressing high levels of the human beta-globin gene. Proc. Natl. Acad. Sci. U. S. A. 92, 6728–6732 (1995).

3. Cong, L. et al. Multiplex genome engineering using CRISPR/Cas systems. Science 339, 819–823 (2013).

4. Jinek, M. et al. RNA-programmed genome editing in human cells. eLife 2, e00471 (2013).

5. Hsu, P. D. et al. DNA targeting specificity of RNA-guided Cas9 nucleases. Nat. Biotechnol. 31, 827–832 (2013).

6. Chaudhari, N., Rickard, A. M., Roy, S., Dröge, P. & Makhija, H. A non-viral genome editing platform for site-specific insertion of large transgenes. Stem Cell Res. Ther. 11, 380 (2020).

7. Pelea, O. et al. Programmable genome editing in human cells using RNA-guided bridge recombinases. Science 391, eadz1884 (2026).

8. Tou, C. J. et al. Immune evasive DNA donors and recombinases license kilobase-scale writing. Nature 1–11 (2026) doi:10.1038/s41586-026-10241-z.

9. Geurts, A. M. et al. Gene transfer into genomes of human cells by the sleeping beauty transposon system. Mol. Ther. J. Am. Soc. Gene Ther. 8, 108–117 (2003).

10. Ivics, Z. et al. Transposon-mediated Genome Manipulations in Vertebrates. Nat. Methods 6, 415–422 (2009).

11. Lampe, G. D. et al. Targeted DNA integration in human cells without double-strand breaks using CRISPR-associated transposases. Nat. Biotechnol. 42, 87–98 (2024).

12. Witte, I. P. et al. Programmable gene insertion in human cells with a laboratory-evolved CRISPR-associated transposase. Science 388, eadt5199 (2025).

13. Fong, J. H. C. & Ceroni, F. Transgene integration in mammalian cells: The tools, the challenges, and the future. Cell Syst. 16, (2025).

14. Blaese, R. M. et al. T lymphocyte-directed gene therapy for ADA-SCID: initial trial results after 4 years. Science 270, 475–480 (1995).

15. Tremblay, J. P., Annoni, A. & Suzuki, M. Three Decades of Clinical Gene Therapy: From Experimental Technologies to Viable Treatments. Mol. Ther. 29, 411–412 (2021).

16. Musunuru, K. et al. Patient-Specific In Vivo Gene Editing to Treat a Rare Genetic Disease. N. Engl. J. Med. 392, 2235–2243 (2025).

17. Roth, T. L. & Marson, A. Genetic Disease and Therapy. Annu. Rev. Pathol. 16, 145–166 (2021).

18. June, C. H. & Sadelain, M. Chimeric Antigen Receptor Therapy. N. Engl. J. Med. 379, 64–73 (2018).

19. Müller, F. et al. CD19 CAR-T cells for treatment-refractory autoimmune diseases: the phase 1/2 CASTLE basket trial. Nat. Med. 32, 1142–1151 (2026).

20. Blaeschke, F. et al. Modular pooled discovery of synthetic knockin sequences to program durable cell therapies. Cell 186, 4216–4234.e33 (2023).

21. Roth, T. L. et al. Non-viral intron knock-ins for targeted gene integration into human T cells and for T-cell selection. Nat. Biomed. Eng. 9, 1309–1319 (2025).

22. Tousley, A. M. et al. Co-opting signalling molecules enables logic-gated control of CAR T cells. Nature 615, 507–516 (2023).

23. Roybal, K. T. et al. Engineering T Cells with Customized Therapeutic Response Programs Using Synthetic Notch Receptors. Cell 167, 419–432.e16 (2016).

24. Foisey, M. G. et al. Synthetic Hybrid Receptors for Safer and Programmable T Cell Therapy. 2026.01.22.701150 Preprint at 10.64898/2026.01.22.701150 (2026).

25. Kumar, M., Keller, B., Makalou, N. & Sutton, R. E. Systematic determination of the packaging limit of lentiviral vectors. Hum. Gene Ther. 12, 1893–1905 (2001).

26. Yacoub, N. al, Romanowska, M., Haritonova, N. & Foerster, J. Optimized production and concentration of lentiviral vectors containing large inserts. J. Gene Med. 9, 579–584 (2007).

27. Rommel, P. C., Engel, N. W., Levine, B. L. & June, C. H. Efficient Production of Chimeric Antigen Receptor (CAR) T Cells with Transgenes Exceeding 10 kb Using Lentiviral Vectors. J. Vis. Exp. JoVE e69121 (2026) doi:10.3791/69121.

28. Cavazzana-Calvo, M. et al. Gene therapy of human severe combined immunodeficiency (SCID)-X1 disease. Science 288, 669–672 (2000).

29. Hacein-Bey-Abina, S. et al. LMO2-associated clonal T cell proliferation in two patients after gene therapy for SCID-X1. Science 302, 415–419 (2003).

30. Kohn, L. A. & Kohn, D. B. Gene Therapies for Primary Immune Deficiencies. Front. Immunol. 12, (2021).

31. Harrison, S. J. et al. CAR+ T-Cell Lymphoma after Cilta-cel Therapy for Relapsed or Refractory Myeloma. N. Engl. J. Med. 392, 677–685 (2025).

32. Aleman, A. et al. Targeted Therapy of CAR+ T-Cell Lymphoma after Anti-BCMA CAR-T. N. Engl. J. Med. 393, 823–825 (2025).

33. He, X., Li, Y.-X. & Feng, B. New Turns for High Efficiency Knock-In of Large DNA in Human Pluripotent Stem Cells. Stem Cells Int. 2018, 9465028 (2018).

34. Blanch-Asensio, A. et al. STRAIGHT-IN enables high-throughput targeting of large DNA payloads in human pluripotent stem cells. Cell Rep. Methods 2, 100300 (2022).

35. Blanch-Asensio, A., Grandela, C., Mummery, C. L. & Davis, R. P. STRAIGHT-IN: a platform for rapidly generating panels of genetically modified human pluripotent stem cell lines. Nat. Protoc. 21, 429–472 (2024).

36. Roth, T. L. et al. Reprogramming human T cell function and specificity with non-viral genome targeting. Nature 559, 405–409 (2018).

37. Wang, Z., Zhou, Z.-J., Liu, D.-P. & Huang, J.-D. Single-stranded oligonucleotide-mediated gene repair in mammalian cells has a mechanism distinct from homologous recombination repair. Biochem. Biophys. Res. Commun. 350, 568–573 (2006).

38. Schumann, K. et al. Generation of knock-in primary human T cells using Cas9 ribonucleoproteins. Proc. Natl. Acad. Sci. 112, 10437–10442 (2015).

39. Eyquem, J. et al. Targeting a CAR to the TRAC locus with CRISPR/Cas9 enhances tumour rejection. Nature 543, 113–117 (2017).

40. Wang, K. et al. Basic enables selection-free efficient knockin of large DNA in primary human T cells. Mol. Ther. 34, 2309–2323 (2026).

41. Oh, S. A. et al. High-efficiency nonviral CRISPR/Cas9-mediated gene editing of human T cells using plasmid donor DNA. J. Exp. Med. 219, e20211530 (2022).

42. Li, H. et al. Design and specificity of long ssDNA donors for CRISPR-based knock-in. 178905 Preprint at 10.1101/178905 (2019).

43. Shy, B. R. et al. High-yield genome engineering in primary cells using a hybrid ssDNA repair template and small-molecule cocktails. Nat. Biotechnol. 41, 521–531 (2023).

44. Zhang, Y. et al. Machine learning-optimized long single-stranded DNA synthesis technology empowers high-precision diagnostic–therapeutic integration in living cells. Nucleic Acids Res. 54, gkag131 (2026).

45. Iyer, S. et al. Efficient Homology-Directed Repair with Circular Single-Stranded DNA Donors. CRISPR J. 5, 685–701 (2022).

46. Letort, G. et al. Circular single stranded DNA potentiates non-viral gene insertion in hematopoietic stem and progenitor cells. Nat. Commun. 16, 10125 (2025).

47. Xie, K. et al. Efficient non-viral immune cell engineering using circular single-stranded DNA-mediated genomic integration. Nat. Biotechnol. 43, 1821–1832 (2025).

48. Velangani, H. G., Ghosh, A., Singh, S. & Kiran, S. Strategies and considerations for the generation of ssDNA-Based HDR templates for CRISPR-based genome editing. BMC Genomics 27, 245 (2026).

49. Durrant, M. G. et al. Systematic discovery of recombinases for efficient integration of large DNA sequences into the human genome. Nat. Biotechnol. 41, 488–499 (2023).

50. Fanton, A. et al. Site-specific DNA insertion into the human genome with engineered recombinases. Nat. Biotechnol. 1–14 (2025) doi:10.1038/s41587-025-02895-3.

51. Anzalone, A. V. et al. Programmable deletion, replacement, integration and inversion of large DNA sequences with twin prime editing. Nat. Biotechnol. 40, 731–740 (2022).

52. Yarnall, M. T. N. et al. Drag-and-drop genome insertion of large sequences without double-strand DNA cleavage using CRISPR-directed integrases. Nat. Biotechnol. 41, 500–512 (2023).

53. Pandey, S. et al. Efficient site-specific integration of large genes in mammalian cells via continuously evolved recombinases and prime editing. Nat. Biomed. Eng. 9, 22–39 (2025).

54. Tatiossian, K. J. et al. Rational Selection of CRISPR-Cas9 Guide RNAs for Homology-Directed Genome Editing. Mol. Ther. J. Am. Soc. Gene Ther. 29, 1057–1069 (2021).

55. Schubert, M. S. et al. Optimized design parameters for CRISPR Cas9 and Cas12a homology-directed repair. Sci. Rep. 11, 19482 (2021).

56. Riesenberg, S. et al. Simultaneous precise editing of multiple genes in human cells. Nucleic Acids Res. 47, e116 (2019).

57. Liu, M. et al. Methodologies for Improving HDR Efficiency. Front. Genet. 9, 691 (2019).

58. Selvaraj, S. et al. High-efficiency transgene integration by homology-directed repair in human primary cells using DNA-PKcs inhibition. Nat. Biotechnol. 42, 731–744 (2024).

59. Yuan, B. et al. Modulation of the microhomology-mediated end joining pathway suppresses large deletions and enhances homology-directed repair following CRISPR-Cas9-induced DNA breaks. BMC Biol. 22, 101 (2024).

60. Kath, J. et al. Pharmacological interventions enhance virus-free generation of TRAC-replaced CAR T cells. Mol. Ther. Methods Clin. Dev. 25, 311–330 (2022).

61. Cullot, G. et al. Genome editing with the HDR-enhancing DNA-PKcs inhibitor AZD7648 causes large-scale genomic alterations. Nat. Biotechnol. 43, 1778–1782 (2025).

62. Aird, E. J., Lovendahl, K. N., St. Martin, A., Harris, R. S. & Gordon, W. R. Increasing Cas9-mediated homology-directed repair efficiency through covalent tethering of DNA repair template. Commun. Biol. 1, 54 (2018).

63. Ling, X. et al. Improving the efficiency of precise genome editing with site-specific Cas9-oligonucleotide conjugates. Sci. Adv. 6, eaaz0051 (2020).

64. Ghasemi, H. I. et al. Interstrand crosslinking of homologous repair template DNA enhances gene editing in human cells. Nat. Biotechnol. 41, 1398–1404 (2023).

65. Jin, Y.-Y. et al. Enhancing homology-directed repair efficiency with HDR-boosting modular ssDNA donor. Nat. Commun. 15, 6843 (2024).

66. Li, W. et al. Efficient high-precision transgene knock-in by Recombinases (Redα/β)-enhanced DNA integration-CRISPR-Cas9 (RED-CRISPR). Nat. Commun. 17, 538 (2025).

67. Riesenberg, S. et al. Efficient high-precision homology-directed repair-dependent genome editing by HDRobust. Nat. Methods 20, 1388–1399 (2023).

68. Perez-Bermejo, J. A. et al. Functional screening in human HSPCs identifies optimized protein-based enhancers of Homology Directed Repair. Nat. Commun. 15, 2625 (2024).

69. Karasu, M. E. et al. Removal of TREX1 activity enhances CRISPR–Cas9-mediated homologous recombination. Nat. Biotechnol. 43, 1168–1176 (2025).

70. Burrell, W. H. et al. Rational design of synthetic proteins using a genome-scale CRISPR screen. 2026.02.19.706875 Preprint at 10.64898/2026.02.19.706875 (2026).

71. Wang, X. et al. A transgene-encoded cell surface polypeptide for selection, in vivo tracking, and ablation of engineered cells. Blood 118, 1255–1263 (2011).

72. Allen, A. G. et al. A highly efficient transgene knock-in technology in clinically relevant cell types. Nat. Biotechnol. 42, 458–469 (2024).

73. Chang, C. R. et al. SEED-Selection enables high-efficiency enrichment of primary T cells edited at multiple loci. Nat. Biotechnol. 43, 2043–2053 (2025).

74. Webber, B. R. et al. Cas9-induced targeted integration of large DNA payloads in primary human T cells via homology-mediated end-joining DNA repair. Nat. Biomed. Eng. 8, 1553–1570 (2024).

75. Tommasi, A. et al. Efficient nonviral integration of large transgenes into human T cells using Cas9-CLIPT. Mol. Ther. Methods Clin. Dev. 33, 101437 (2025).

76. Nguyen, D. N. et al. Polymer-stabilized Cas9 nanoparticles and modified repair templates increase genome editing efficiency. Nat. Biotechnol. 38, 44–49 (2020).

77. Kornete, M., Marone, R. & Jeker, L. T. Highly Efficient and Versatile Plasmid-Based Gene Editing in Primary T Cells. J. Immunol. 200, 2489–2501 (2018).

78. Yao, X. et al. Homology-mediated end joining-based targeted integration using CRISPR/Cas9. Cell Res. 27, 801–814 (2017).

79. Jing, R. et al. Cas9-Cleavage Sequences in Size-Reduced Plasmids Enhance Nonviral Genome Targeting of CARs in Primary Human T Cells. Small Methods 5, 2100071 (2021).

80. Allen, G. M. et al. Synthetic cytokine circuits that drive T cells into immune-excluded tumors. Science 378, eaba1624 (2022).

81. Garcia, J. et al. Naturally occurring T cell mutations enhance engineered T cell therapies. Nature 626, 626–634 (2024).

82. Lesueur, L. L., Mir, L. M. & André, F. M. Overcoming the Specific Toxicity of Large Plasmids Electrotransfer in Primary Cells *In Vitro*. Mol. Ther. - Nucleic Acids 5, e291 (2016).

83. Søndergaard, J. N. et al. Successful delivery of large-size CRISPR/Cas9 vectors in hard-to-transfect human cells using small plasmids. Commun. Biol. 3, 319 (2020).

84. Suzuki, K. et al. In vivo genome editing via CRISPR/Cas9 mediated homology-independent targeted integration. Nature 540, 144–149 (2016).

85. Balke-Want, H. et al. Homology-independent targeted insertion (HITI) enables guided CAR knock-in and efficient clinical scale CAR-T cell manufacturing. Mol. Cancer 22, 100 (2023).

86. Staal, J., Alci, K., De Schamphelaire, W., Vanhoucke, M. & Beyaert, R. Engineering a minimal cloning vector from a pUC18 plasmid backbone with an extended multiple cloning site. BioTechniques 66, 254–259 (2019).

87. Chen, Z.-Y., He, C.-Y., Ehrhardt, A. & Kay, M. A. Minicircle DNA vectors devoid of bacterial DNA result in persistent and high-level transgene expression in vivo. Mol. Ther. J. Am. Soc. Gene Ther. 8, 495–500 (2003).

88. Oliynyk, R. T., Mahas, A., Karpinski, E. & Church, G. M. Plasmid2MC: efficient cell-free generation of high-purity minicircle DNA for genome editing in mammalian cells. Commun. Biol. 8, 1778 (2025).

89. Chanani, P., Rezaei, N., Dormiani, K., Shokatian, M. & Ata-abadi, N. S. Progress and prospect of minicircle as a minimized non-viral DNA vector in gene therapy and regenerative medicine. Mol. Ther. Nucleic Acids 36, 102682 (2025).

90. Krall, W. J. et al. Increased levels of spliced RNA account for augmented expression from the MFG retroviral vector in hematopoietic cells. Gene Ther. 3, 37–48 (1996).

91. Tomberg, K. et al. Intronization enhances expression of S-protein and other transgenes challenged by cryptic splicing. 2021.09.15.460454 Preprint at 10.1101/2021.09.15.460454 (2021).

92. DeKelver, R. C. et al. Functional genomics, proteomics, and regulatory DNA analysis in isogenic settings using zinc finger nuclease-driven transgenesis into a safe harbor locus in the human genome. Genome Res. 20, 1133–1142 (2010).

93. Pantazis, C. B. et al. A reference human induced pluripotent stem cell line for large-scale collaborative studies. Cell Stem Cell 29, 1685–1702.e22 (2022).

94. Straathof, K. C. et al. An inducible caspase 9 safety switch for T-cell therapy. Blood 105, 4247–4254 (2005).

